# Modulation of photosystem II heterogeneity and stomatal functioning contribute to sustaining photochemistry and photoprotection in mangroves exposed to a cold wave

**DOI:** 10.1101/2024.04.08.588527

**Authors:** John Sunoj. V. Sebastian, Sonal Mathur, Nabil. I. Elsheery, Li Yan, Hans Lambers, Aidan. W. Short, Alison. K.S. Wee, Anjana Jajoo, Amy Ny Aina Aritsara, Tadashi Kajita, Kun-Fang Cao

**Author notes:** Both authors contributed equally to the study (John Sunoj. V. Sebastian, Sonal Mathur). Authors for correspondence: Kun-fang Cao, State Key Laboratory of Conservation and Utilization of Subtropical Agro-bio-resources and Guangxi Key Laboratory of Forest Ecology and Conservation, College of Forestry, Guangxi University, Nanning 530004, Guangxi, PR China. Current address: Institute of Soil Water and Environmental Sciences, Volcani Center, 68 HaMaccabim Road, P.O.B 15159, Rishon LeZion, 7505101, Israel.

## Abstract

Cold waves restrict the distribution of mangroves. This study examined the contribution of PSII heterogeneity and stomatal functioning to sustaining photochemistry and photoprotection in mangroves during a cold wave. We exposed eight populations of Kandelia obovata (cold-tolerant) and, Bruguiera gymnorhiza (cold-susceptible) from different latitudes to 27/20°C (favorable) and 10/3°C (chilling; simulated cold wave) day and night temperatures. Multiple trait responses imply that cold waves affected K. obovata the least. Significant changes in chlorophyll fluorescence transients (photosystem II [PSII]) with a slight decrease in the redox status of P700 (photosystem I [PSI]) imply a greater impact of a cold wave on PSII. During the cold wave, photochemical efficiency of PSII, efficiency of the water-splitting complex, light absorptance, stomatal pore area, cyclic electron flow, nonphotochemical quenching, and number of active PSIIα and PSII QB reducing centers decreased, while light transmittance, night respiration, and inactive PSII QB nonreducing, PSIIβ, and γ centers increased in both species. The population of K. obovata from the coldest latitudinal site (Fujian, China) was least affected by cold wave due to local evolutionary adaptations. Modulation of PSII heterogeneity and stomatal functioning is important to sustaining photochemistry and photoprotection in mangroves to cope with cold waves.

**Highlights:** PSII heterogeneity and stomatal functioning support mangroves to cope with cold waves.

Local evolutionary adaptations promote the cold tolerance of mangrove populations.

## Introduction

Mangroves are one of the biosphere’s highly productive ecosystems; however, mangrove species found in tropical and subtropical intertidal zones are poorly adapted to cold (Coldren & Proffitt, 2017; Saenger et al., 2019). In marginal distribution areas, occasional cold waves (rapid drop in temperature [0-10°C] for short periods) during winter cause severe damage to mangroves (Zheng *et al*., 2016; Guo et al., 2018;) and restrain their distribution and poleward migration under global warming (Cavanaugh et al., 2014; Madrid et al., 2014; Yan et al., 2021).

Foliar gas exchange (photosynthesis and respiration) is a vital physiological process for growth and development of plants, and one of the primary targets of abiotic stress factors (e.g., high temperature, cold, drought, flooding, and salinity) (Sunoj et al., 2017; Elsheery et al., 2020; Maswada et al., 2020; Yan et al., 2021). These stress factors, including different forms of cold stress, severely inhibit and/or block photosynthesis and modify respiration which interrupts the carbon balance (carbon trade-off [photosynthesis vs. respiration]) (Sunoj et al., 2016, 2022; Yan et al., 2021). Likewise, studies reported analogous impacts of cold on the foliar gas exchange of mangroves (Chen et al., 2017; Yan et al., 2021). In general, under favorable day and night temperatures, a major share of incident light on leaves absorbed by the light-harvesting antenna complex of photosystems (PSII and PSI) is utilized for (1) photochemistry by transferring the energy to reaction centers (photochemical quenching [qP]; light-dependent reactions to generate ATP and NADPH, which are utilized for CO_2_-assimilation reactions to fix CO_2_ by converting it into carbohydrates) and excess light energy is converted to (2) fluorescence and (3) dissipated as heat (non-photochemical quenching; qE [NPQ]; xanthophyll cycle). These three energy-utilization processes occur in a balance (Muller et al., 2001; Saez et al., 2019; Miyake, 2020; Sunoj et al., 2022).

Cold adversely affects stomatal and nonstomatal controls over foliar gas exchange which are associated with several interrelated metabolic processes of qP and qE causing an imbalance between the above energy-utilization processes (Allen & Ort, 2001; Knight et al., 2004; Saez et al., 2019). As a consequence, photoinhibition of PSII and PSI occurs and gradually accumulates reactive oxygen species (ROS; superoxide [O2^•–^], hydrogen peroxide [H_2_O_2_], singlet oxygen [^1^O_2_] and hydroxyl radicals [^•^OH]), which results in ROS-mediated oxidative damage to DNA, proteins, and lipids which leads to disruption of plant growth (Derks et al., 2015; Takagi et al., 2016; Elsheery et al., 2020). On the other hand, during cold, excess incident light further intensifies photoinhibition and photodamage due to prolonged excitation of PSII and PSI reaction centers (Elsheery et al. 2008; Huang et al. 2017; Zheng et al. 2016; Yang et al. 2017; Sunoj et al., 2022). Under such hostile circumstances, efficient management of incident light and utilization of energy from absorbed light by the light-dependent reactions along with efficient CO_2_-assimilation reactions mitigates the intensity of stress impacts to some extent, and such capability accounts for the magnitude of stress tolerance (Sunoj et al., 2023).

To manage energy from excess incident light and prevent photoinhibition of photosystems and photooxidative damages under abiotic stresses, multiple photoprotection mechanisms are activated before the surplus production of injurious ROS (Miyake, 2010; Belgio et al., 2014; Neto et al., 2017). Rapidly-acting photoprotection mechanisms and supporting physiological-biochemical modulations, in addition to morphological changes, involve (a) the stomatal level regulation over stomatal functioning (speed of stomatal closure and opening and stomatal pore area) (Raven, 2014; Jurczyk et al., 2019) and (b) at photosystem level including PSII heterogeneity (antenna-size and reducing-side) (Bukhov & Carpentier, 2000; Belgio et al., 2014; Mathur et al., 2021), moderate photoinhibition of PSII (Sunoj et al., 2022), photorespiration, night respiration (Yan et al., 2021), cyclic electron flow around PSI (CEF), non-photochemical quenching (NPQ; xanthophyll cycle) and alternative electron flow (water-water cycle [WWC] or the Mehler–ascorbate peroxidase pathway [MAP]) (Miyake, 2010; Neto et al., 2017). However, the extent of activation of different photoprotection mechanisms in response to diverse stresses varies with the magnitude of stress tolerance or vice versa across plant species which can be genetic and irreversible within their lifespan and/or involve phenotypic plasticity (reversible) according to the existing microclimatic conditions (Körner, 2016).

There are a few studies conducted that demonstrate interspecific variation in the magnitude of reduction in photosynthesis, and activation of diverse photoprotection mechanisms in mangroves under cold (Zheng et al., 2016, 2016a; Chen et al., 2017; Song et al., 2020; Yan et al., 2021). For mangroves, we recently reported that along with multiple other photoprotection mechanisms, night respiration takes part in protecting thylakoid membrane damage to minimize the reduction of phytochemical efficiency and photoinhibition under cold (Yan et al., 2021). Different studies concluded that night respiration plays a vital role in sustaining various photoprotection mechanisms and maintaining carbon balance by utilizing assimilated CO_2_ (photosynthates) for growth and repairing photooxidative stress-induced damages in different plant species under diverse stress conditions (Sunoj et al., 2016, 2020; Impa et al., 2018; Mathur et al., 2021). Likewise, proposed roles of PSII heterogeneity (Mathur et al., 2021) and stomatal functioning (Raven, 2014), which have strong control over photosynthesis, contributing to photoprotection and their influence on carbon balance in other plant species have not been well-documented in mangroves during a cold wave.

The reorganization of PSII heterogeneity is an important strategy in plants to overcome various abiotic stresses (Mehta et al., 2010; Mathur et al., 2011a; Tomar et al., 2012). Variability in PSII structure and function of plants is known as PSII heterogeneity. There are two types of PSII heterogeneity, antenna-size and reducing-side heterogeneity. PSII heterogeneity is closely connected with electron transfer activities on the reducing and oxidizing sides of PSII. PSII heterogeneity has been extensively studied; however, how the heterogeneity of PSII changes in response to cold has been studied less. Recently, we compared the PSII heterogeneity of sugarcane cultivars with different levels of cold tolerance under cold conditions revealing a clear genotypic difference (Mathur et al., 2021). The results suggested that PSII heterogeneity can be a photoprotection mechanism that participates in the management of absorption of incident light by altering the optical properties of leaves and regulating the electron transport in response to cold to protect against photoinhibition.

Stomata respond to the changes in microclimatic conditions based on vapor pressure deficit (VPD) and play an important role in gas exchange, carboxylation efficiency, controlling transpirational water loss, stomatal conductance (CO_2_ assimilation), water-use efficiency and nutrient assimilation which in turn influence growth and development (Allen & Ort, 2001; Raven, 2014; Jurczyk et al., 2019; Qie et al., 2024). Hence, stomatal functioning is crucial under stress conditions and is of great interest but not yet fully understood (Drake et al., 2013; Raven, 2014; Jurczyk et al., 2019). However, stomatal functioning protects PSII from photodamage by avoiding xylem failure by limiting transpiration and maintaining photosynthesis (Raven, 2014). Stomata in the majority of the mangrove species are slightly sunken within the epidermis of the abaxial side of the leaves. Such anatomy increases humidity around the stomatal pore and decreases the leaf-to-air VPD and transpiration (Roth-Nebelsick, 2007; Reef & Lovelock, 2015).

Several studies documented the physiological-biochemical responses in mangroves to cold (Zheng et al., 2016; Chen et al., 2017; Yan et al., 2021), but only a few studies have been conducted to understand the effect of cold waves on mangroves (Peereman et al., 2021; He et al., 2023). Unlike other cold stresses, cold waves are expected to have different magnitudes of impacts on plant responses because of the sudden drops in day and night temperatures. In mangroves, there are no studies that document the contribution of modulations of stomatal functioning and PSII heterogeneity to sustaining photochemistry and photoprotection during a cold wave. Hence, to fill this knowledge gap, we monitored responses and correlations among stomatal functioning, PSII heterogeneity, leaf optical properties, photochemistry, and night respiration. We examined the effect of a simulated cold wave on interspecies adaptive variation in responses of mangroves by including populations of two sympatric mangrove species, with different levels of cold tolerance, from different latitudinal locations (Table 1 and Fig 1).

**Table 1:** Geographic coordinates and locations, annual mean temperature, mean daily high and low and average temperatures of coldest quarter of a year at particular geographic location and abbreviations for different populations of the two mangrove species of Rhizophoraceae used in the current study. The climatic data were collected from China Metrological Data Service Center (http://data.cma.cn.html) and Japan Meteorological Agency (https://www.jma.go.jp).

| Species (S)/ populations (P) | Geographical coordinates and locations | Annual mean temperature (°C) | Mean daily high, low and average temperatures (°C) of coldest quarter of a year | Abbreviations for the species and populations |
| --- | --- | --- | --- | --- |
| *Bruguiera gymnorhiza* (L.) Lam |  |  |  | BG |
| S1P1 | Guangxi, China<br>21°29'52.5"N, 109°45'40.0"E | 22.9 | 20.6/ 13.8/ 17.2 | BGG |
| S1P2 | Okinawa, Japan<br>26°39'19.3"N, 128°00'39.0"E | 22.8 | 20.2/ 14.2/ 17.3 | BGO |
| S1P3 | Iriomote, Japan<br>24°24'19.2"N, 123°46'37.1"E | 23.7 | 21.5/ 16.8/ 19.2 | BGI |
| S1P4 | Hainan, China<br>19°37'31.4"N, 110°50'18.5"E | 24.3 | 24.0/ 17.6/ 20.8 | BGH |
| *Kandelia obovata* (S.L.) Yong |  |  |  | KO |
| S2P1 | Guangxi, China<br>21°25'12.4"N, 109°12'44.9"E | 22.9 | 20.3/ 13.8/ 17.0 | KOG |
| S2P2 | Okinawa, Japan<br>26°27'27.7"N, 127°56'22.8"E | 22.4 | 19.7/ 13.2/ 16.4 | KOO |
| S2P3 | Iriomote, Japan<br>24°23'28.2"N, 123°49'26.6"E | 23.7 | 22.1/ 16.6/ 19.4 | KOI |
| S2P4 | Fujian, China<br>23°55'09.5"N, 117°25'14.3"E | 21.3 | 19.8/ 11.1/ 15.9 | KOF |

**Fig. 1:**
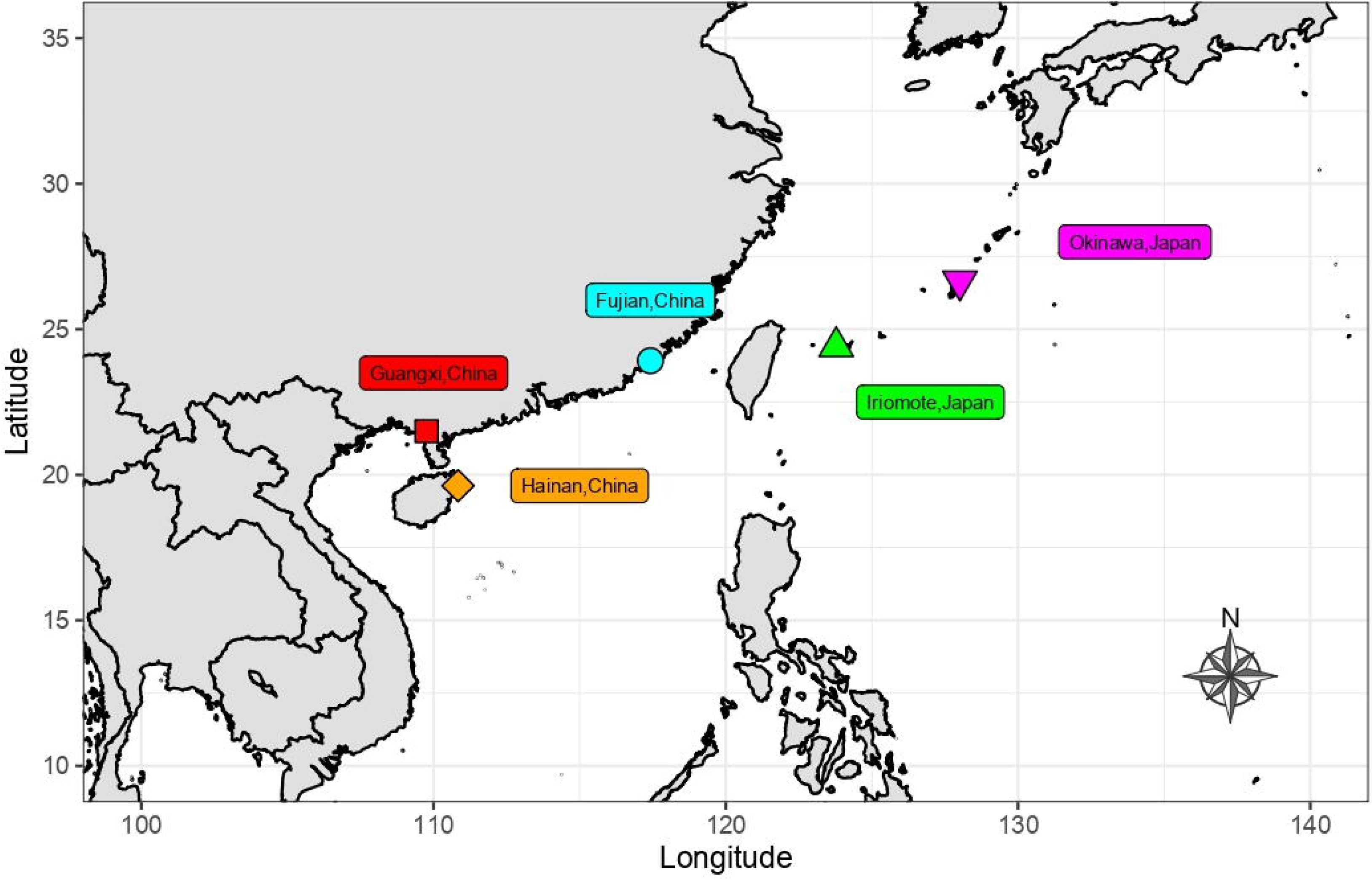
Geographic locations where propagules were sampled (Guangxi and Hainan [China] and Iriomote and Okinawa [Japan]) marked in different shapes and colors.

Specifically, we address the following questions: (1) What are the effects of cold wave on stomatal functioning, PSII heterogeneity, photochemistry, and photoprotection of populations of two mangrove species differing in cold tolerance? (2) How do stomatal functioning and PSII heterogeneity contribute to sustaining photochemistry and photoprotection of populations of two diversely cold-tolerant mangrove species during a cold wave? (3) Are there any correlations between stomatal functioning, PSII heterogeneity, leaf optical properties, photochemistry, photoprotection, and night respiration during a cold wave that contribute to cold tolerance? We tested the hypothesis that photosystems and light-independent reactions of photosynthesis of mangrove populations from colder sites are less affected during a cold wave due to a different response to incident light and efficient utilization of energy by the maintenance of photochemistry by sustaining photoprotection with the support of modulation in stomatal functioning and PSII heterogeneity, and faster night respiration and other photoprotection mechanisms.

## Material and methods

### Plant material and growth condition

The experiment was carried out at Guangxi University, Nanning, China (22.83°N, 108.28°E). Propagules of four populations from each of two mangrove species, *Bruguiera gymnorhiza* (L.) Lam (cold-susceptible) and *Kandelia obovata* (S.L.) Yong (cold-tolerant) (Chen et al., 2017), distributed at sites of different latitudes of China and Japan (Table 1 and Fig. 1) were collected, grown in pots placed in eight plastic tanks for two years and maintained in a glasshouse under favorable growth conditions (mean annual day/night time temperature of 28°C [±8°C]/23°C [±6°C], a photoperiod of 12 h between 06.00 to 18.00 h and mean annual relative humidity (RH) of 77.5% [±17%]). The tanks were uniformly filled with a mixture of NaCl (0.2 M) and half-strength Hoagland’s solutions (Hoagland & Arnon, 1950). Adequate care was taken to maintain a uniform salinity by monitoring the electrical conductivity of the solution in all tanks throughout the experimental period.

To start the experiment, six morphologically similar saplings per population of the two mangrove species were transferred to a walk-in growth chamber (Conviron Model CMP 6050; Winnipeg, MB, Canada) and allowed to acclimate for three days with set microclimatic conditions inside the growth chamber. The set growth chamber conditions were 27/20°C (favorable day and night temperatures [FDNT]; mean daily temperature of 23.5°C, diurnal temperature amplitude of 7°C), 60% of RH, and 12 h photoperiod [06:00 h to 18:00 h]) with photosynthetically active radiation (PAR) of 500 µmol m^-2^ s^-1^ at plant canopy level provided by cool fluorescent lamps. A transition time of 5 h from maximum day (from 10..00 to 17.00 h) to minimum night temperatures (from 22.00 to 05.00 h) and vice versa was followed to replicate the diurnal temperature fluctuation of natural conditions. The maximum photochemical efficiency (F_v_/F_m_) of samplings was measured from physiologically mature leaves using a portable chlorophyll fluorometer (PAM-2100; Walz, Effeltrich, Germany) every day after transfer to the growth chamber to ensure acclimation of plants to set FDNT.

### Cold wave simulation and exposure to cold wave

After acclimation to FDNT, growth chamber temperatures were decreased from 20°C (favorable night temperature) to 10°C (cold day temperature) within a period of 10 h (from 2400 to 1000 h), and on the same day temperature was further decreased to 3°C (cold night temperature) within a period of 5 h (from 17.00 to 22.00 hours) to simulate the occurrence of a cold wave (CDNT). The mean day temperature during the CDNT (10/3°C; cold day/ night temperatures) was 6.5°C and the diurnal temperature amplitude was 7°C and saplings were exposed to CDNT for three days. A common set of physiological measurements in the current study were made on the 3^rd^ day of exposure to FDNT and CDNT.

### Chlorophyll *a* (Chl *a*) fluorescence transients, redox state of P700, and cyclic electron flow (CEF) in PSI

Chlorophyll *a* (Chl *a*) fluorescence transients and OJIP curves were made using a Plant Efficiency Analyzer (Handy PEA; Hansatech Norfolk, England, UK). Measurements were taken from the middle of fully-mature leaves after 30 minutes of dark adjustment (Mathur et al., 2021). The maximum photochemical efficiency of PSII (F_v_/F_m_) was calculated using the following equation:

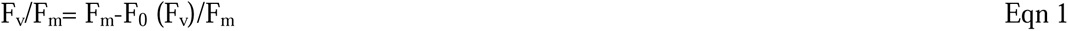

F_m_ refers to the dark-adjusted maximum fluorescence, F_0_ is the dark-adjusted minimum fluorescence and F_v_ is the variable fluorescence.

The efficiency of the water-splitting complex was determined by the ratio of F_v_ and F_0_ (F_v_/F_0_) (Mathur et al., 2011a), and thylakoid membrane damage was calculated from the ratio of F_0_ and F_m_ (F_0_/F_m_) (Sunoj et al., 2017) (Supplementary Table 1).

Performance Index [PI_(abs)_] represents the whole primary photochemical reactions and PI_(abs)_ was calculated by the following equation:

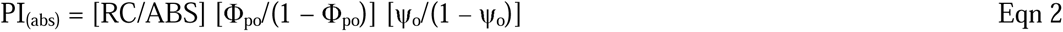

RC/ABS refers to the density of active PSII reaction centers per chlorophyll, Φ_po_/(1−Φ_po_) describes the performance of the light reaction and ψ_o_/(1−ψ_o_) explains the performance of the dark reaction (Mathur et al., 2011a) (Supplementary Table 1). (RC, ABS, Φ_po_ and ψ_o_ represent the reaction centers, absorption, maximum efficiency of PSII, and the probability that an electron can move further than Q_A─_, respectively).

Energy pipeline leaf models were analyzed using Biolyzer HP3 software (Flouomatic lab, MediaSoft, Nyon, Switzerland). The energy pipeline leaf model represents the flow of energy in PSII (Mathur & Jajoo, 2014). ABS/CSo is absorption per cross section (CS), ETo/CSo is electron transport at time zero, TRo/CSo is trapping at time zero per CS and DIo/CSo is dissipation at time zero per CS (Supplementary Table 1).

Light-adjusted Chl *a* fluorescence transients and P700 redox state were recorded using Dual PAM-100 (Heinz Walz, Effeltrich, Germany). Before the measurements, leaves were kept under light (500 µmol m^-2^ s^-1^ PAR) for 30 min; and then the light-adjusted chlorophyll fluorescence and P700 transients were recorded after 3 minutes of exposure to light intensity of 1000 µmol m^-2^ s^-1^ by placing the leaf in the measuring head of the Dual PAM-100. The light-adjusted chlorophyll fluorescence transients were calculated by using the following equations (Genty et al., 1989; Oxborough & Baker, 1997):

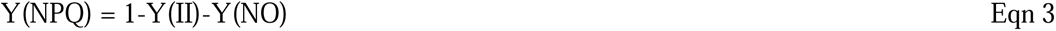

Y(NPQ) is the fraction of energy dissipated as heat through regulated non-photochemical quenching (NPQ), Y(II) is the effective photochemical quantum yield of PSII, and Y(NO) is the quantum yield of non-regulated energy dissipation (Kramer et al., 2004).

Cyclic electron flow (CEF) was calculated (Miyake et al., 2005, 2005a; Sunoj et al., 2022) using the following equation:

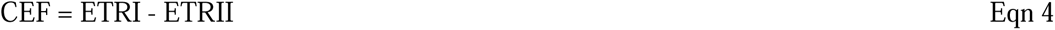

ETRII and ETRI are the electron transport rates of PSII and PSI, respectively.

The maximum photo-oxidizable P700 in PSI (P_m_) was determined by applying a saturation pulse of 10,000 µmol m^-2^ s^-1^ for 300 ms after 30 min of dark adjustment and then applying far-red light for 10 s. The *P*_m_ represents the maximal change of the P700 signal from the fully-reduced state P700 (minimum signal level; P700*) to the fully-oxidized state P700 (maximum signal level; P700^+^) upon application of the saturation pulse.

### PSII Heterogeneity

#### Reducing-side heterogeneity

Reducing-side heterogeneity (Q_B_-reducing and Q_B_-non-reducing centers) was calculated using the double hit (pulse) method following Strasser and Tsimilli-Michael (1998) and Mathur et al. (2011).

#### Antenna-size heterogeneity

Antenna-size heterogeneity was calculated using the DCMU [3-(3,4-dichlorophenyl)-1,1-dimethylurea] poisoning method, in terms of the percentage of PSII alpha (α), beta (β) and gamma (γ) centers following Hsu et al. (1989) and Mathur et al. (2011).

#### Night respiration

Leaf night respiration was recorded using a portable photosynthesis system (Li-6400XT; LI-COR, NE, USA) from the fully-mature leaves between 22,00–23,00 h after four hours of exposure to darkness (0 µ mol m^-2^ s^-1^ PAR; 17.00–22.00 h) on the 3^rd^ day of exposure to FDNT and CDNT. The block temperature inside the leaf chamber of the photosynthesis system was set to the respective night temperature of each temperature condition (20°C [FDNT] and 3°C [CDNT; cold wave]), while the CO_2_ concentration (400 µmol mol^-1^) and flow rate (100 µmol s^-1^) were adjusted for minimum fluctuation (Sunoj et al., 2016, 2020).

#### Stomatal morphology and functioning and leaf temperature

Morphological traits of the stomata (stomatal pore length, width, and area) were measured following Muralikrishna et al., 2010. Microscopic images were captured using a digital camera (Leica MC190 HD, Heerbrugg, Switzerland) mounted on a light microscope (Leica DN 3000LED, Wetzlar, Germany) at a magnification of 40x. The polygon selection tool of imageJ software (Fiji Version 1.52n, NIH, USA) was used to measure stomatal morphology. Stomatal functioning was explained in terms of changes in stomatal pore area. The leaf temperature was recorded using an infrared thermal camera (FLIR C2 Compact Thermal Camera; FLIR Systems Inc. Oregon, USA).

#### Leaf optical properties and chlorophyll index

Leaf optical properties (light reflectance, transmittance, and absorptance) were measured between 1100 h to 1200 h using a miniature leaf spectrometer (CI-710; CID Bio-science, Camas, WA, USA). Chlorophyll index was recorded using a chlorophyll meter (SPAD-502 Plus; Konica Minolta Inc., Japan).

#### Statistical and data analysis

For statistical analyses, measured data of different traits from six biological replicates per population of each species under FDNT and CDNT were used. The statistical design was a split plot. Analysis of variance (ANOVA) was performed using a generalized linear model (GLM) in SPSS (SPSS Inc. Ver.16, USA) to test the significance of differences in traits between the populations of the two species (Table 2). Student’s t-test was used to compare the response of the two species (irrespective of populations) under FDNT and CDNT (Supplementary Tables 2 to 4). Pearson correlation analysis was performed for all the traits separately for FDNT and CDNT and in combination (irrespective of population; Supplementary Tables 5 to 7). The means of individual traits of different populations of each species under FDNT and CDNT were compared using Duncan’s multiple range test (DMRT; Supplementary Tables 2 to 4). Statistical analyses and graphs of Chl *a* fluorescence transients and the heterogeneity of PSII were performed and generated using Origin Pro8 and GraphPad Prism 5.01(GraphPad Software, Inc.; La Jolla, CA, USA), respectively.

**Table 2:**
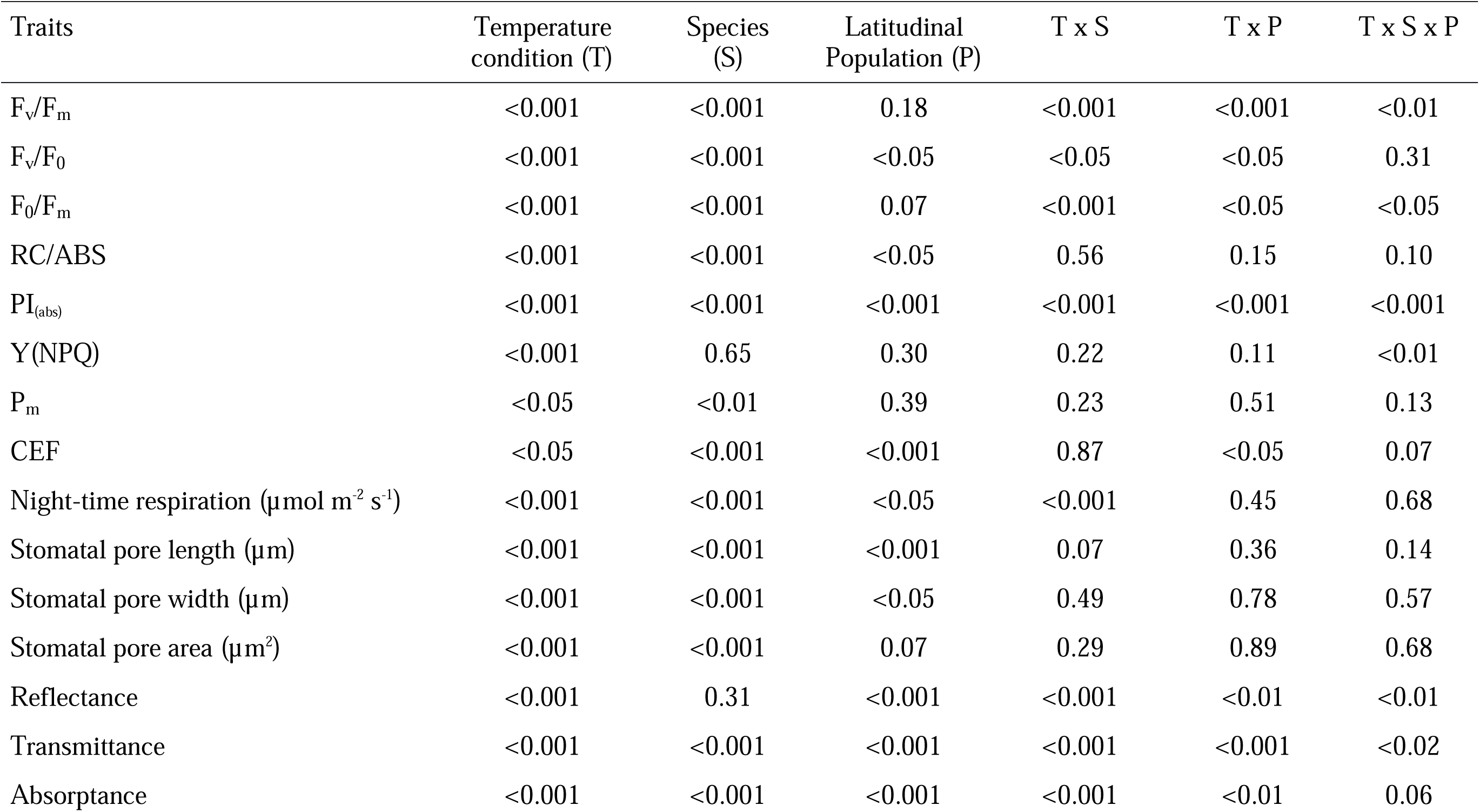
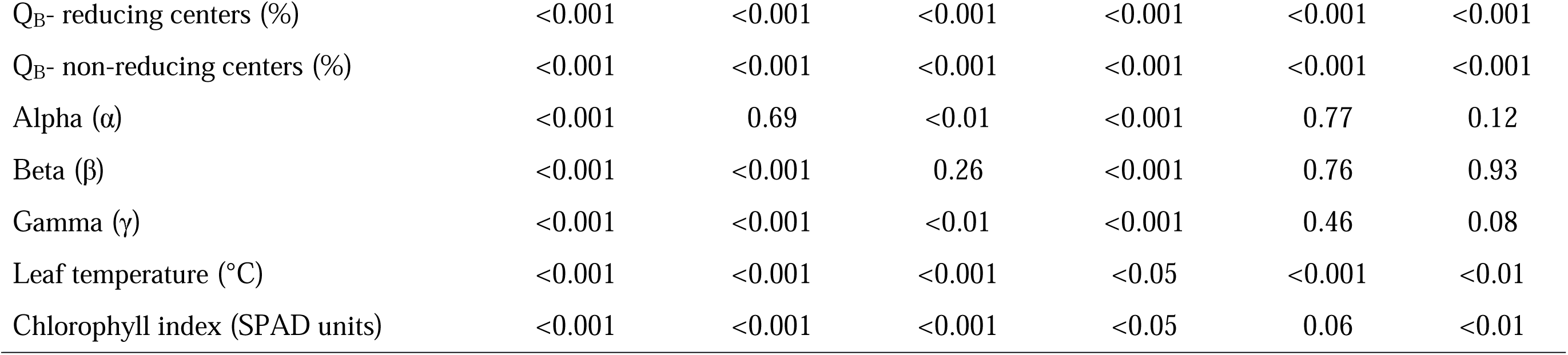
Probability values of the effect of different temperature conditions (T), species (S), latitudinal population (P) and their interaction (T x S x P) on chlorophyll *a* (Chl*a*) fluorescence transients, redox state of P700, cyclic electron flow in PSI, night respiration, stomatal morphology and functioning, leaf optical properties, PSII heterogeneity, leaf temperature and chlorophyll index. F_v_/F_m_: maximum quantum yield of PSII in the dark-adjusted state (relative units); F_v_/F_0_: status of water-splitting complex (relative units), F_0_/F_m_: thylakoid membrane damage (relative units); RC/ABS: active PSII reaction centers per chlorophyll; PI_(abs)_:performance index; Y(NPQ): effective quantum yield of NPQ; P_m_: maximum photo-oxidizable P700 (ΔI/I); CEF: cyclic electron flow.

## Results

We observed significant (*P*<0.001) effects of a cold wave across all the studied traits, except for P_m_ and CEF (<0.05) of mangrove species. Out of 22 recorded traits, three between species and six among populations showed no significant difference. At the same time, the interaction effect of temperature conditions (favorable [FDNT] and cold [CDNT; cold wave] day and night temperatures), species, and populations were significant (*P*<0.05) on 10 traits (Table 2).

### Effect of cold wave on chlorophyll *a* fluorescence

Regardless of species and population, significant decreases in F_v_/F_m_, F_v_/F_0_, RC/ABS, Y(NPQ) and PI_(abs)_ and increases in F_0_/F_m_ were during CDNT and *Kandelia obovata* showed the least changes (F_v_/F_m_ [46%], F_v_/F_0_ [82%], RC/ABS [55%], Y(NPQ) [29%], PI_(abs)_ [95%] and F_o_/F_m_ [260%]). At the population level, KOF and BGO displayed the least changes in the above traits (Table 3 and Supplementary Table 2), except for Y(NPQ).

**Table 3:** Percentage change in chlorophyll *a* (Chl*a*) fluorescence transients, redox state of P700, cyclic electron flow in PSI, night respiration, stomatal morphology and functioning, leaf optical properties, PSII heterogeneity, leaf temperature and chlorophyll index of populations of *Bruguiera gymnorhiza* (BG) and *Kandelia obovata* (KO) from favorable (FDNT) to cold (CDNT; cold wave) day and night temperature conditions. BGG: BG from Guangxi, China, BGH: BG from Hainan, China, BGI: BG from Iriomote, Japan, BGO: BG from Okinawa, Japan, KOG: KO from Guangxi, China, KOF: KO from Fujian, China, KOI: KO from Iriomote, Japan and KOO: KO from Okinawa, Japan. F_v_/F_m_: maximum quantum yield of PSII in the dark-adjusted state (relative units); F_v_/F_0_: status of water-splitting complex (relative units), F_0_/F_m_: thylakoid membrane damage (relative units); RC/ABS: active PSII reaction centers per chlorophyll; PI_(abs)_:performance index; Y(NPQ): effective quantum yield of NPQ; P_m_: maximum photo-oxidizable P700 (ΔI/I); CEF: cyclic electron flow.

| Species/ Populations | *Bruguiera gymnorhiza* |  |  |  |  | *Kandelia obovata* |  |  |  |  |
| --- | --- | --- | --- | --- | --- | --- | --- | --- | --- | --- |
|  | BGG | BGH | BGI | BGO | Mean BG populations | KOG | KOF | KOI | KOO | Mean KO population |
| Traits | Percentage change (%) |  |  |  |  |  |  |  |  |  |
| $F_v/F_m$ | -82 | -84 | -67 | -59 | -73 | -49 | -38 | -52 | -46 | -46 |
| $F_v/F_0$ | -96 | -96 | -92 | -91 | -94 | -81 | -76 | -87 | -83 | -82 |
| $F_0/F_m$ | 342 | 314 | 329 | 247 | 311 | 235 | 207 | 321 | 267 | 260 |
| RC/ABS | -83 | -83 | -74 | -61 | -75 | -57 | -46 | -61 | -60 | -55 |
| $PI_{(abs)}$ | -99 | -99 | -99 | -97 | -99 | -96 | -92 | -97 | -96 | -95 |
| Y(NPQ) | -25 | -45 | -34 | -53 | -38 | -42 | 8 | -68 | -13 | -29 |
| $P_m$ | -3 | 2 | -20 | -8 | -8 | -2 | -4 | -11 | -11 | -7 |
| CEF | -0.2 | -9 | -31 | -29 | -18 | -7 | 61 | -20 | -20 | -4 |
| Night respiration ($\mu\text{mol m}^{-2} \text{s}^{-1}$) | 53 | 53 | 85 | 66 | 63 | 67 | 80 | 121 | 86 | 87 |
| Stomatal pore length ($\mu\text{m}$) | -3 | -7 | -6 | -9 | -6 | -4 | -3 | -7 | -5 | -3 |
| Stomatal pore width ($\mu\text{m}$) | -47 | -32 | -51 | -49 | -46 | -45 | -13 | -32 | -48 | -40 |
| Stomatal pore area ($\mu\text{m}^2$) | -47 | -28 | -49 | -44 | -44 | -28 | -26 | -44 | -50 | -34 |
| Reflectance | -18 | 0 | -7 | -33 | -15 | 7 | 14 | -7 | 0 | 4 |
| Transmittance | 30 | 18 | 50 | 79 | 44 | 8 | -9 | 23 | 15 | 8 |
| Absorptance | -5 | -6 | -12 | -8 | -8 | -3 | -1 | -3 | -3 | -3 |
| Q<sub>B</sub>-reducing centers (%) | -36 | -39 | -20 | -20 | -29 | -23 | -21 | -31 | -23 | -26 |
| Q<sub>B</sub>-nonreducing centers (%) | 40 | 46 | 31 | 30 | 37 | 35 | 33 | 56 | 36 | 38 |
| Alpha ($\alpha$) | -14 | -13 | -18 | -17 | -16 | -29 | -21 | -30 | -22 | -26 |
| Beta ($\beta$) | 11 | 15 | 22 | 14 | 16 | 13 | 3 | 13 | 16 | 13 |
| Gamma ($\gamma$) | 27 | 15 | 7 | 33 | 21 | 100 | 122 | 118 | 62 | 100 |
| Leaf temperature (°C) | -52 | -60 | -50 | -60 | -55 | -54 | -59 | -52 | -53 | -55 |
| Chlorophyll index (SPAD units) | -6 | -4 | -2 | -2 | -4 | -2 | 2 | 2 | -6 | -2 |

Chlorophyll *a* (chl *a*) fluorescence transient curves (OJIP) of *Bruguiera gymnorhiza* and *K. obovata* under FDNT and cold CDNT showed typical OJIP polyphasic curves (Fig. 2). A sharp decline in OJIP curves was observed in both *B. gymnorhiza* and *K. obovata* during CDNT with prominent damping in J, I and P phases. Among all the *K. obovata* populations, KOF was the least affected by the CDNT, while among the *B. gymnorhiza* populations, BGO was the least affected (Fig.2).

**Fig. 2:**
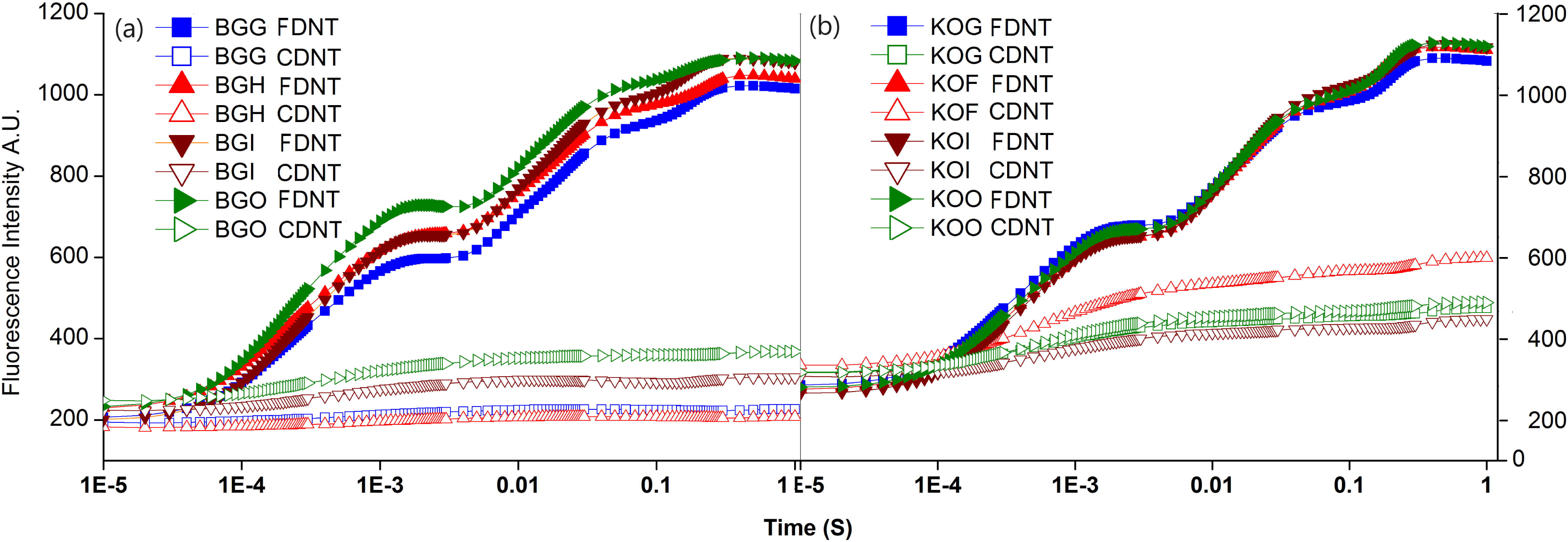
Chlorophyll (Chl *a*) fluorescence transient (OJIP) curve of populations of (A) *Bruguiera gymnorhiza* (BG) and (B) *Kandelia obovata* (KO) under favorable (FDNT) and cold (CDNT [cold wave]) day and night temperature conditions. BGG: BG from Guangxi, China, BGH: BG from Hainan, China, BGI: BG from Iriomote, Japan, BGO: BG from Okinawa, Japan, KOG: KO from Guangxi, China, KOF: KO from Fujian, China, KOI: KO from Iriomote, Japan and KOO: KO from Okinawa, Japan.

The chl *a* fluorescence analysis based on the cross-section (CS) of the energy pipeline leaf model confirmed that in both species CDNT led to a greater reduction in electron transport (ETo/CSo) and trapping per cross-section (TRo/CSo) followed by a gradual reduction in absorption (ABS/CSo), while there was an increase in dissipation (DIo/CSo) (Fig. 3 [representative figures of KOF and BGO based on energy pipeline model]). The CDNT increased the number of inactive reaction centers in both species as indicated by the filled circles in the output images of the energy pipeline leaf model and the overall, least change in model outputs found in *K. obovata* (Fig. 3).

**Fig. 3:**
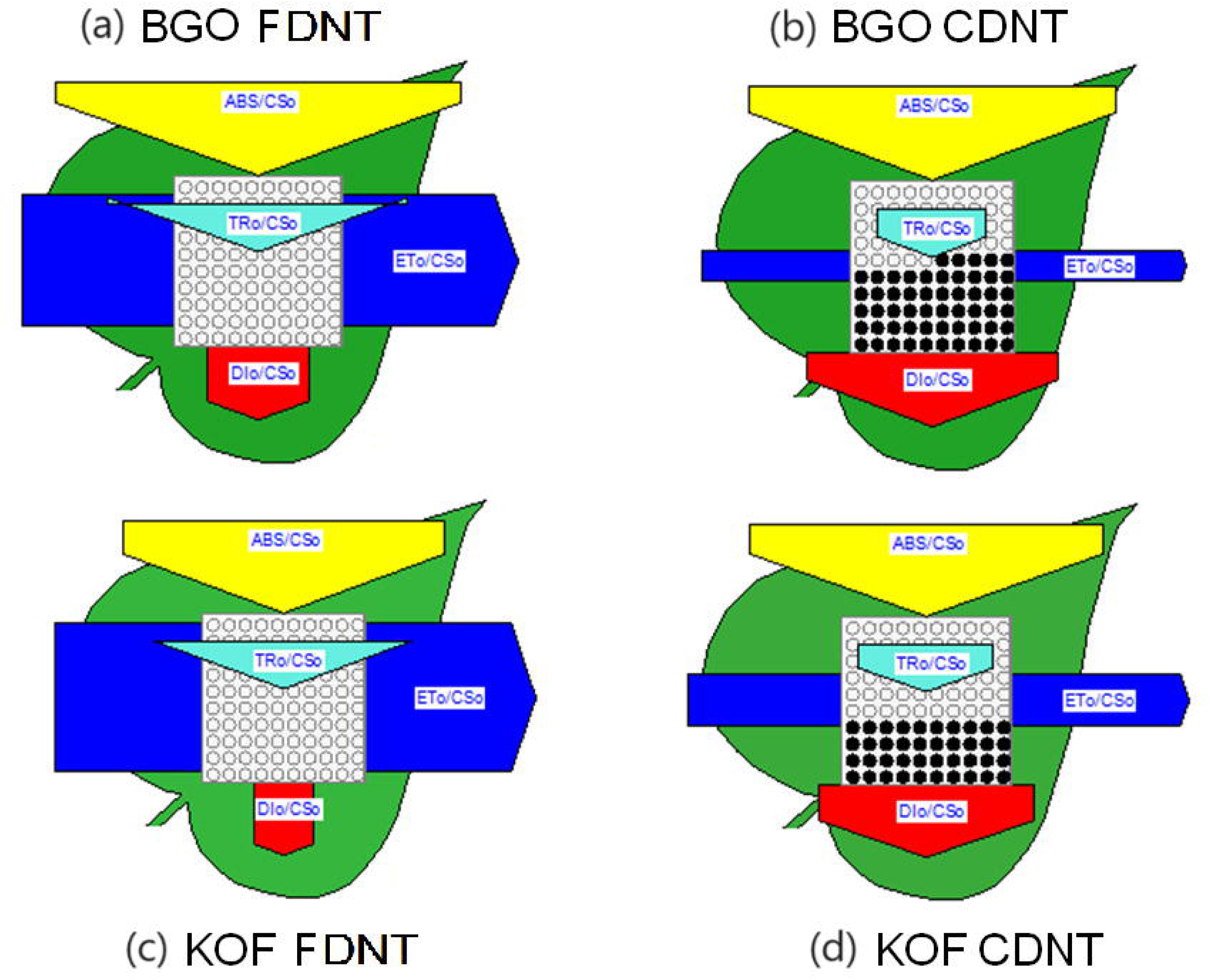
Representative energy pipeline leaf model of light energy utilization of PSII under favorable (FDNT) [(A) *Bruguiera gymnorhiza* (BG) and (C) *Kandelia obovata* (KO)] and cold (CDNT [cold wave]) [(B) BG and (D) KO] day and night temperature conditions. Where A and B represents BGO: BG from Okinawa, Japan and C and D represents KOF: KO from Fujian, China. The open and filled circles indicate the open and closed reaction centers of PSII, respectively. ABS/CSo: absorption per cross section (CS), ETo/CSo: electron transport at time zero, TRo/CSo: trapping at time zero per CS and DIo/CSo: dissipation at time zero per CS.

### Effect of a cold wave on P700 redox state, cyclic electron flow (CEF), and night respiration

A significant decrease in P_m_ was observed across the populations of *B. gymnorhiza* (8%) and *K. obovata* (7%), except for BGH, which showed an increase of 2%. Conversely, CEF and Y(NPQ) increased by 61 and 8% in KOF, respectively, but decreased in all other populations of *B. gymnorhiza* and *K. obovata* (Table 3 and Supplementary Table 2). Night respiration significantly increased during CDNT and was fastest in *K. obovata* (87%). Among the different populations, the increase in night respiration was greatest in KOI (121%), but the actual values were high in KOF and BGO (Table 3 and Supplementary Table 2).

### Effect of a cold wave on PSII heterogeneity, leaf optical properties, and chlorophyll index

Irrespective of species and population, PSII antenna-size heterogeneity, percentage contribution of PSII_α_ centers (active centers) significantly decreased, while the percentage contribution of the less active PSII_β_ and PSII_γ_ centers increased during CDNT (Fig. 4). At the same time, the percentage of PSII_β_ centers was greater in *B. gymnorhiza*, while the percentage of PSIIγ centers was greater in *K. obovata* (Table 3, Fig. 4 and Supplementary Table 3). Among the *B. gymnorhiza* populations, the smallest decrease (13%) in PSII_α_ centers was in BGH, while the greatest increase in PSII_β_ and PSII_γ_ centers was observed in BGI (22%) and BGO (33%), respectively (Table 3). Conversely, the smallest reduction (21%) in PSII_α_ centers, the greatest increase in PSII_β_ (16%) was for KOO and the greatest increase (122%) in PSII_γ_ centers was in KOF (Table 3).

**Fig. 4:**
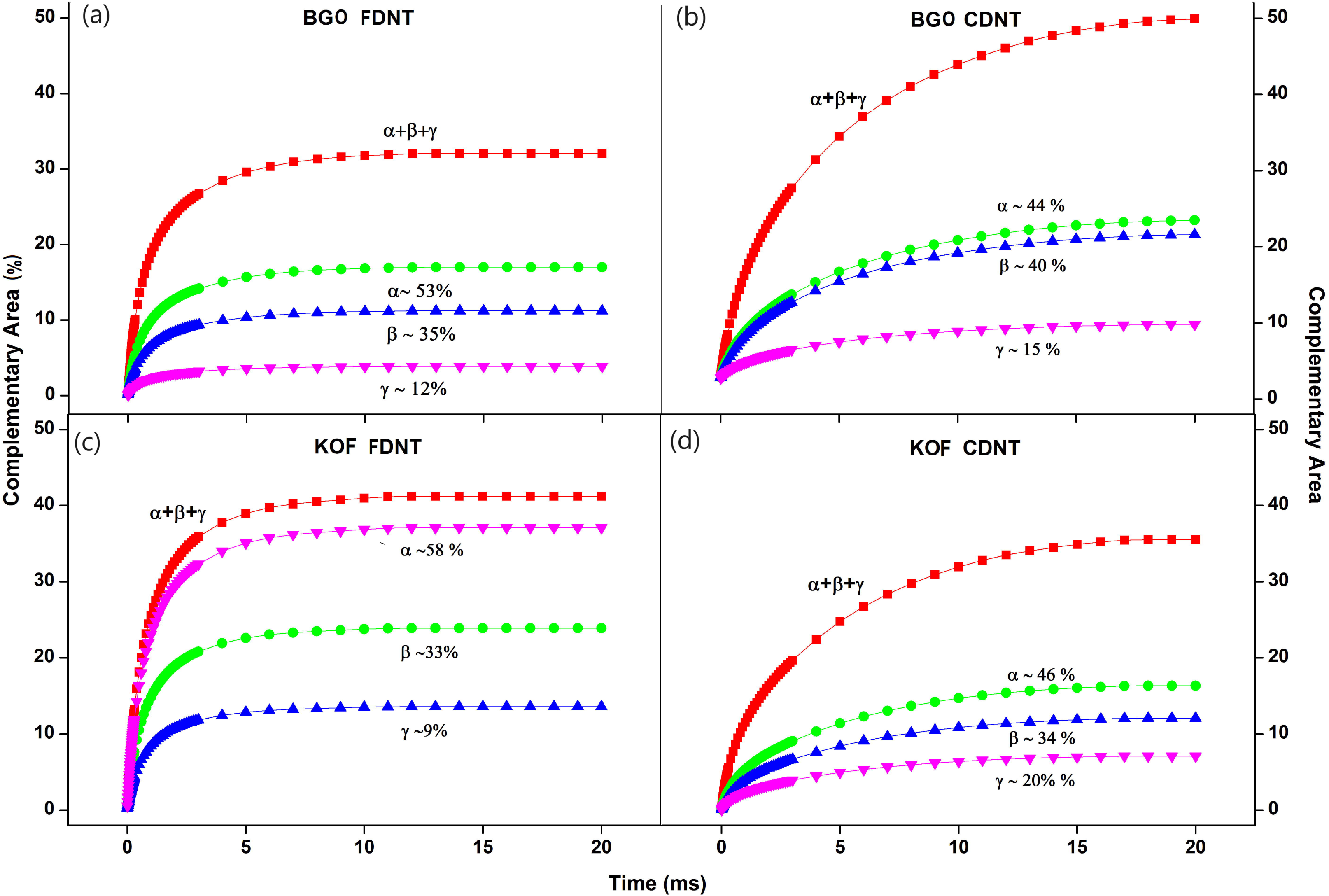
Representative complementary area growth curve showing percentage of PSIIα, PSIIβ and PSIIγ centers of PSII under favorable (FDNT) [(A) *Bruguiera gymnorhiza* (BG) and (C) *Kandelia obovata* (KO)] and cold (CDNT [cold wave]) [(B) BG and (D) KO] day and night temperature conditions. Where A and B represent BGO: BG from Okinawa, Japan, and C and D represent KOF: KO from Fujian, China.

During CDNT, in PSII reducing-side heterogeneity, Q_B_-reducing centers significantly (P<0.001) decreased (*B. gymnorhiza* [29%] and in *K. obovata* [26%]) and the Q_B_-nonreducing centers increased (*B. gymnorhiza* [37%] and *K. obovata* [38%]; Table 3 and Supplementary Table 3). Among the populations, BGO and BGI showed the least decrease (20%) in Q_B_-reducing centers followed by KOF (21%) (Table 3). Simultaneously, the smallest increase in Q_B_-nonreducing centers was observed in BGO (30%) and KOF (33%) (Table 3).

Leaf optical properties of the two species responded differentially during CDNT. The transmittance increased in both species, while absorptance decreased. Simultaneously, reflectance showed a mixed response. The *K. obovata* displayed the least increase in transmittance (8%) and decrease in absorptance (3%) (Table 3 and Supplementary Table 3). The greatest decrease in reflectance (33%) and increase in transmittance (79%) was observed in BGO, while the smallest decrease in absorptance (1%) was observed in KOF (Table 3 and Supplementary Table 3).

Regardless of population, CDNT caused a slight decrease of 4 and 2% in chlorophyll index in *B. gymnorhiza* and *K. obovata*, respectively (Table 3). Among the populations, the greatest reduction in chlorophyll index was observed in KOO and BGG (6%) (Table 3 and Supplementary Table 3).

### Effect of the cold wave on stomatal functioning and leaf temperature

We found greater decreases in stomatal pore length (6%), width (46%) and area (44%) in *B. gymnorhiza* during CDNT (Table 3, Supplementary Fig. 1 and Supplementary Table 4). Among the populations, we observed the smallest decrease in all stomatal traits in KOF. The smallest decrease in stomatal pore area was in KOF (26%) and BGH (28%) (Table 3, Supplementary Fig. 1 and Supplementary Table 4). Leaf temperature decreased in both species and the least in BOI (50%) and KOI (52%) (Table 3). On average, the reduction in leaf temperature was the same in *B. gymnorhiza* and *K. obovata* (Table 6). Meanwhile, the actual values showed low leaf temperature in KOF and BGO (Supplementary Table 4 and Supplementary Fig. 2).

### Effect of a cold wave on the correlation between traits

Regardless of temperature conditions, species, and populations, we found significant correlations among the traits (Supplementary Table 5). The correlations were stable or varied when analyzed separately for FDNT and CDNT (Supplementary Tables 6 and 7). To focus on and emphasize alteration in PSII heterogeneity and stomatal functioning, and its influence on carbon balance and photoprotection under FDNT and CDNT, the most significant and relevant correlations were selected and presented in graphical format (Figs 5 to 10).

**Fig. 5:**
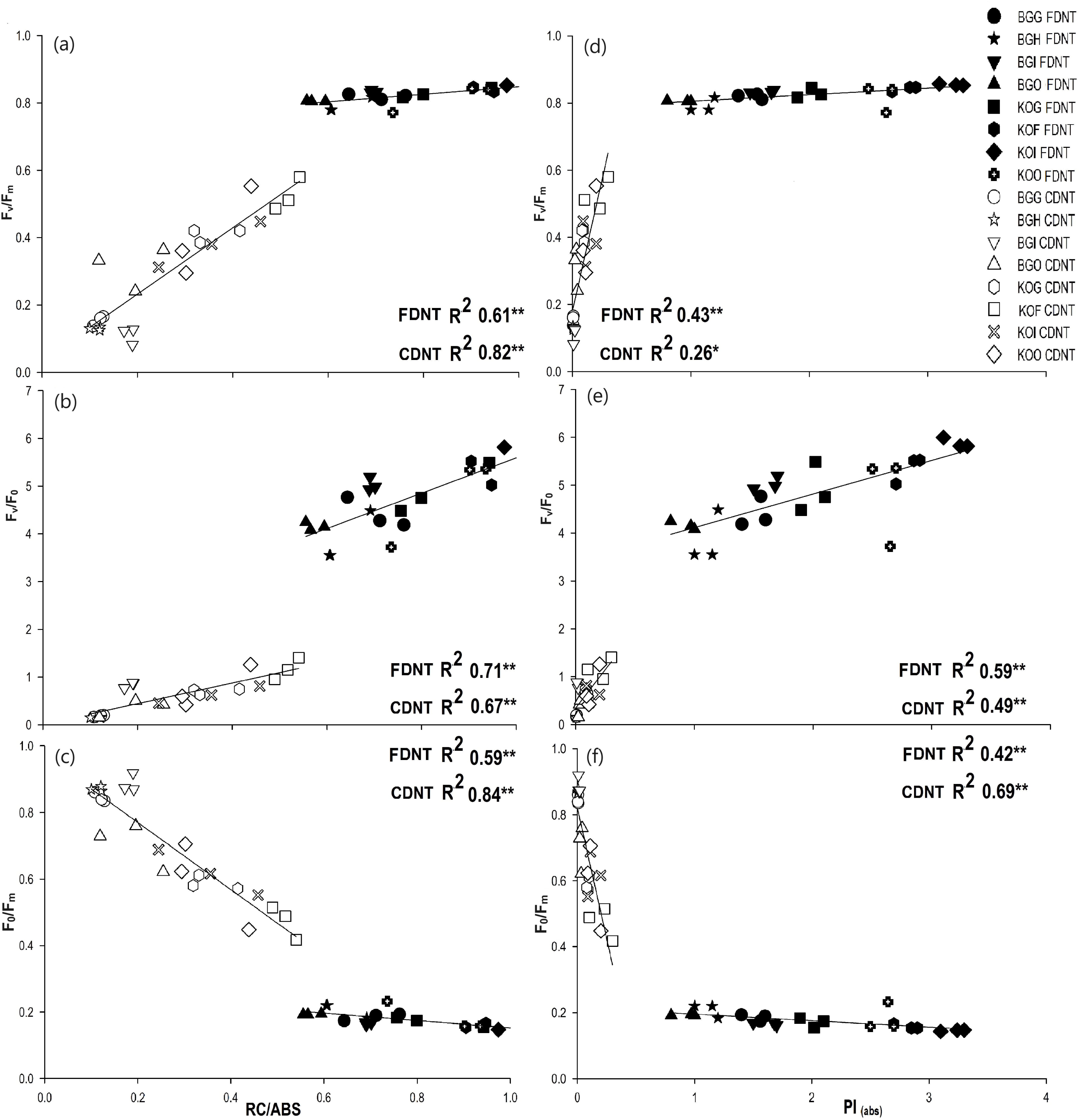
Correlation of RC/ABS and PI_(abs)_ with (A and D) F_v_/F_m_, (B and E) F_v_/F_0_ and (C and F) F_m_/F_0_ of different populations of *Bruguiera gymnorhiza* (BG) and *Kandelia obovata* (KO) under favorable (FDNT; *closed symbols*) and cold (CDNT [cold wave]; *open symbols*) day and night temperature conditions. Coefficient of determination (R^2^) followed by * and ** corresponding to significance at *P*<0.05 and *P*<0.01, respectively. BGG: BG from Guangxi, China, BGH: BG from Hainan, China, BGI: BG from Iriomote, Japan, BGO: BG from Okinawa, Japan, KOG: KO from Guangxi, China, KOF: KO from Fujian, China, KOI: KO from Iriomote, Japan and KOO: KO from Okinawa, Japan.

**Fig. 6:**
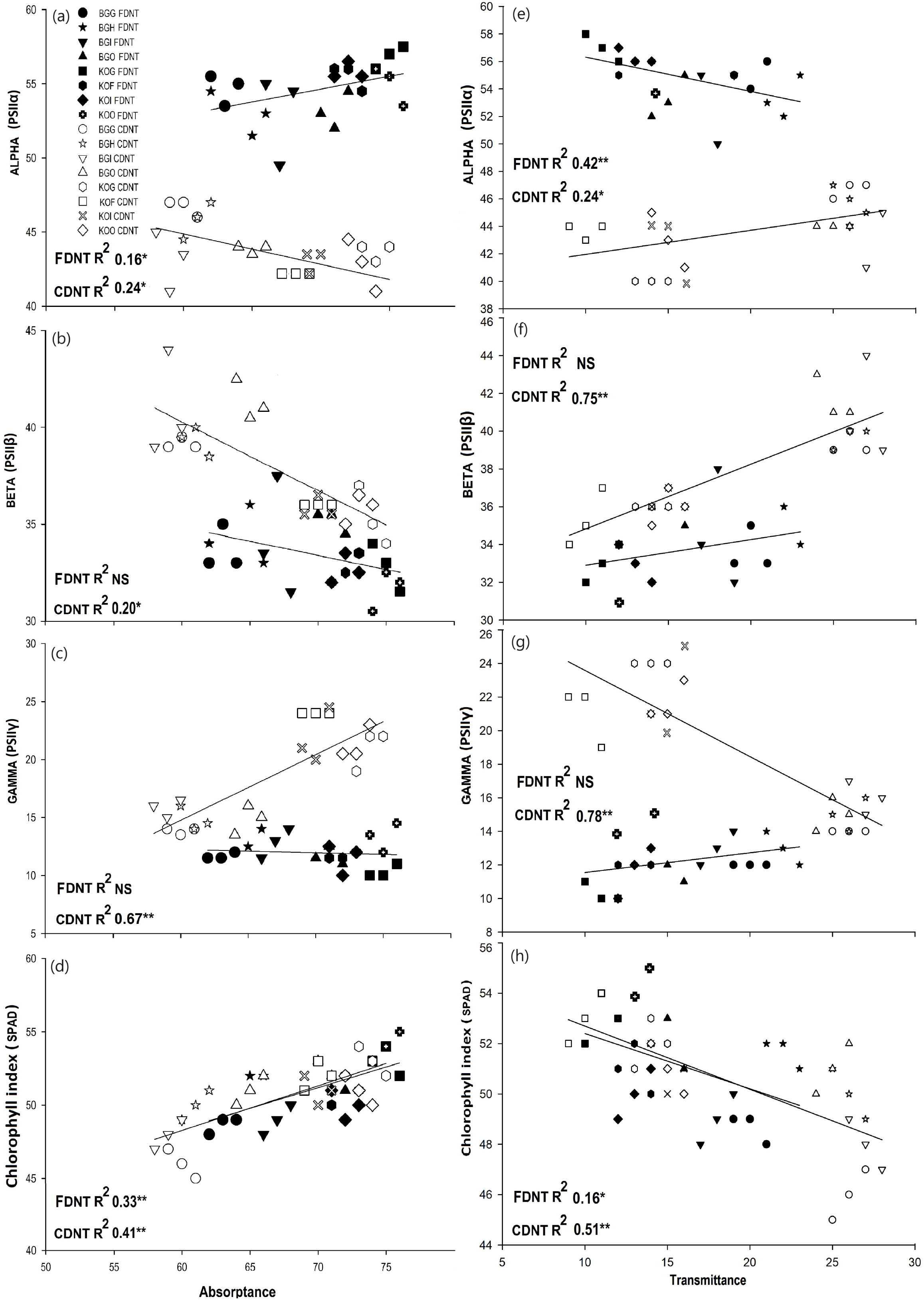
Correlation of leaf absorptance and transmittance with antenna-size heterogeneity of PSII [(A) and (D) alpha (PSIIα), (B) and (E) beta (PSIIβ) and (C) and (F) gamma (PSIIγ)] and (D) and (G) chlorophyll index of different populations of *Bruguiera gymnorhiza* (BG) and *Kandelia obovata* (KO) under favorable (FDNT; *closed symbols*) and cold (CDNT [cold wave]; *open symbols*) day and night temperature conditions. Coefficient of determination (R^2^) followed by NS, * and ** corresponding to non-significance, significance at *P*<0.05 and *P*<0.01, respectively. BGG: BG from Guangxi, China, BGH: BG from Hainan, China, BGI: BG from Iriomote, Japan, BGO: BG from Okinawa, Japan, KOG: KO from Guangxi, China, KOF: KO from Fujian, China, KOI: KO from Iriomote, Japan and KOO: KO from Okinawa, Japan.

**Fig. 7:**
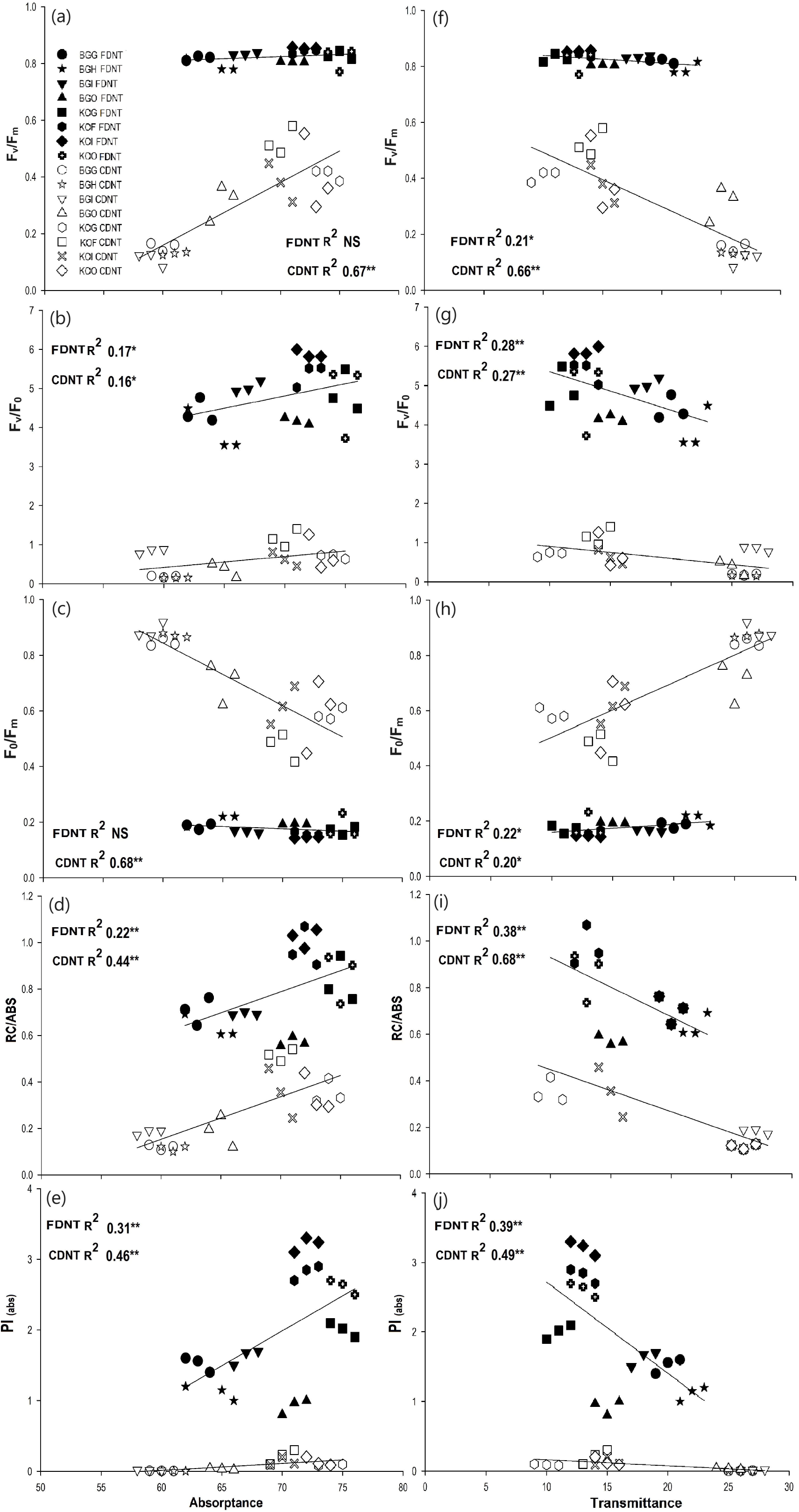
Correlation of leaf absorptance and transmittance with (A and F) F_v_/F_m_, (B and G) F_v_/F_0_, (C and H) F_m_/F_0_, (D and I) RC/ABS and (D and J) PI_(abs)_ of different populations of *Bruguiera gymnorhiza* (BG) and *Kandelia obovata* (KO) under favorable (FDNT; *closed symbols*) and cold (CDNT [cold wave]; *open symbols*) day and night temperature conditions. Coefficient of determination (R^2^) followed by NS, * and ** corresponding to non-significance, significance at *P*<0.05 and *P*<0.01, respectively. BGG: BG from Guangxi, China, BGH: BG from Hainan, China, BGI: BG from Iriomote, Japan, BGO: BG from Okinawa, Japan, KOG: KO from Guangxi, China, KOF: KO from Fujian, China, KOI: KO from Iriomote, Japan and KOO: KO from Okinawa, Japan.

**Fig. 8:**
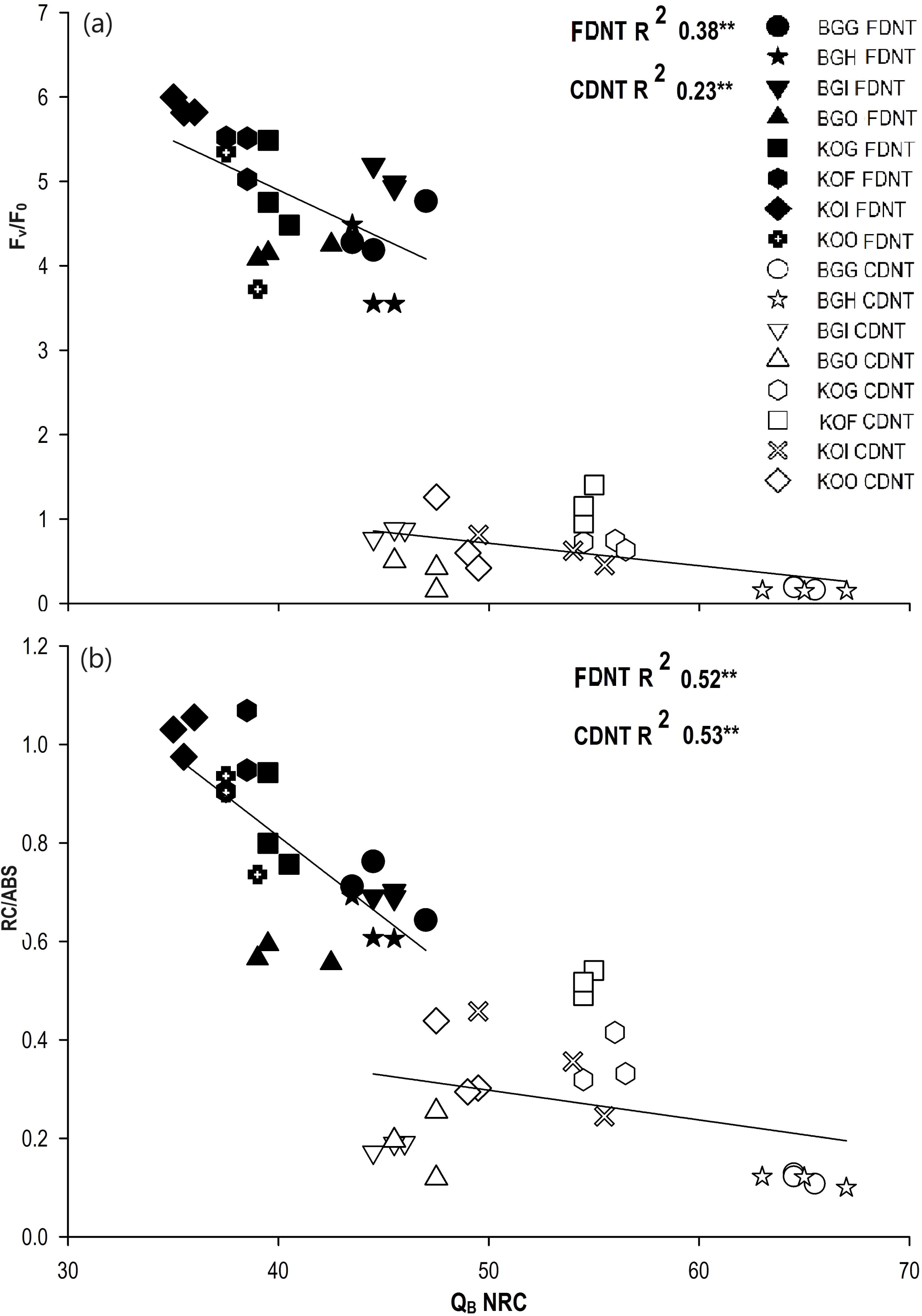
Correlation of Q_B_ non-reducing centers (Q_B_-non-reducing centers) with (A) F_v_/F_m_, and (B) RC/ABS of different populations of *Bruguiera gymnorhiza* (BG) and *Kandelia obovata* (KO) under favorable (FDNT; *closed symbols*) and cold (CDNT [cold wave]; *open symbols*) day and night temperature conditions. Coefficient of determination (R^2^) followed by ** corresponding to significance at *P*<0.01. BGG: BG from Guangxi, China, BGH: BG from Hainan, China, BGI: BG from Iromote, Japan, BGO: BG from Okinawa, Japan, KOG: KO from Guangxi, China, KOF: KO from Fujian, China, KOI: KO from Iriomote, Japan and KOO: KO from Okinawa, Japan.

**Fig. 9:**
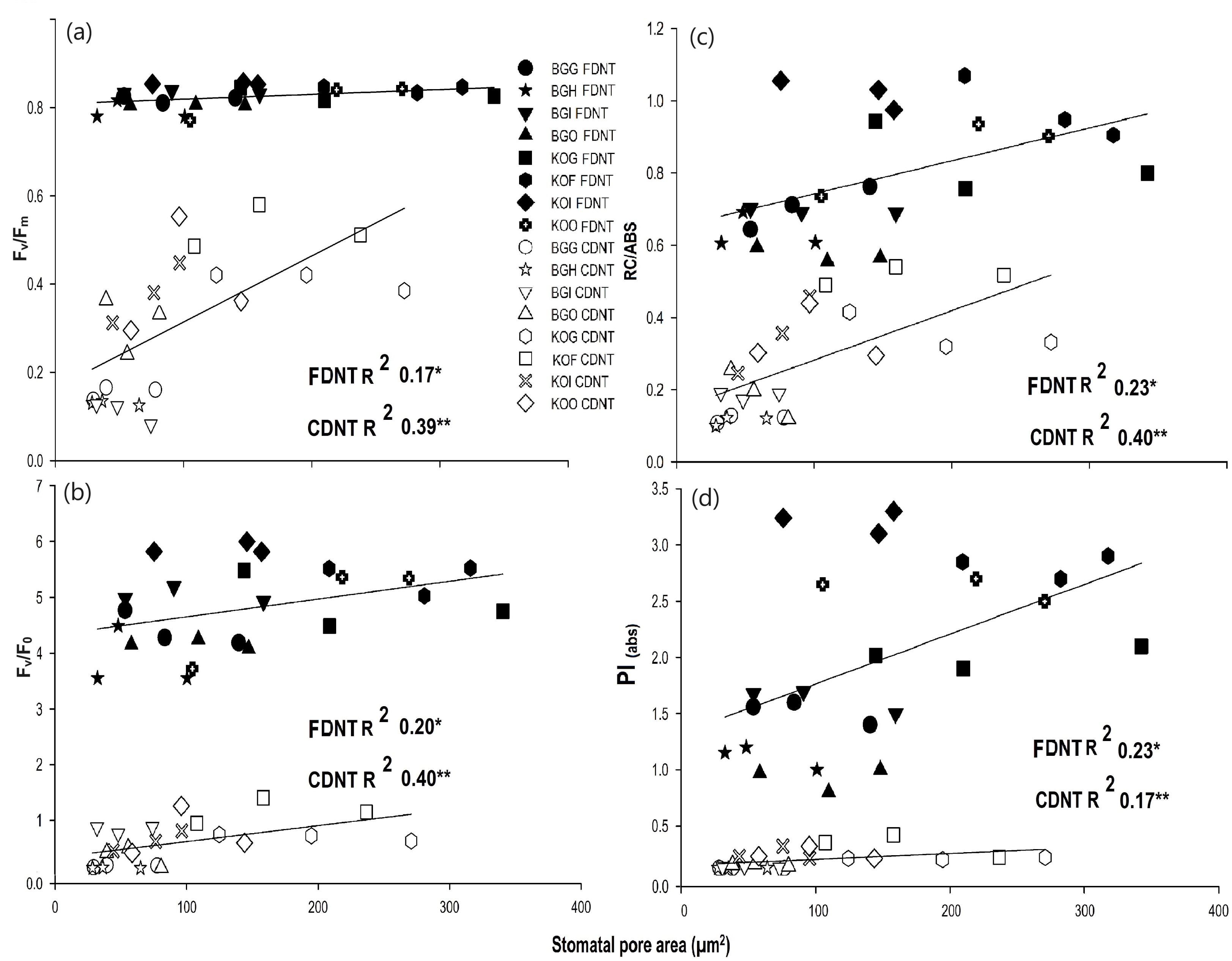
Correlation of stomatal pore area with (A) F_v_/F_m_, (B) F_v_/F_0_, (C) RC/ABS and (D) PI_(abs)_ of different populations of *Bruguiera gymnorhiza* (BG) and *Kandelia obovata* (KO) under favorable (FDNT; *closed symbols*) and cold (CDNT [cold wave]; *open symbols*) day and night temperature conditions. Coefficient of determination (R^2^) followed by * and ** corresponding to significance at *P*<0.05 and *P*<0.01, respectively. BGG: BG from Guangxi, China, BGH: BG from Hainan, China, BGI: BG from Iriomote, Japan, BGO: BG from Okinawa, Japan, KOG: KO from Guangxi, China, KOF: KO from Fujian, China, KOI: KO from Iriomote, Japan and KOO: KO from Okinawa, Japan.

**Fig. 10:**
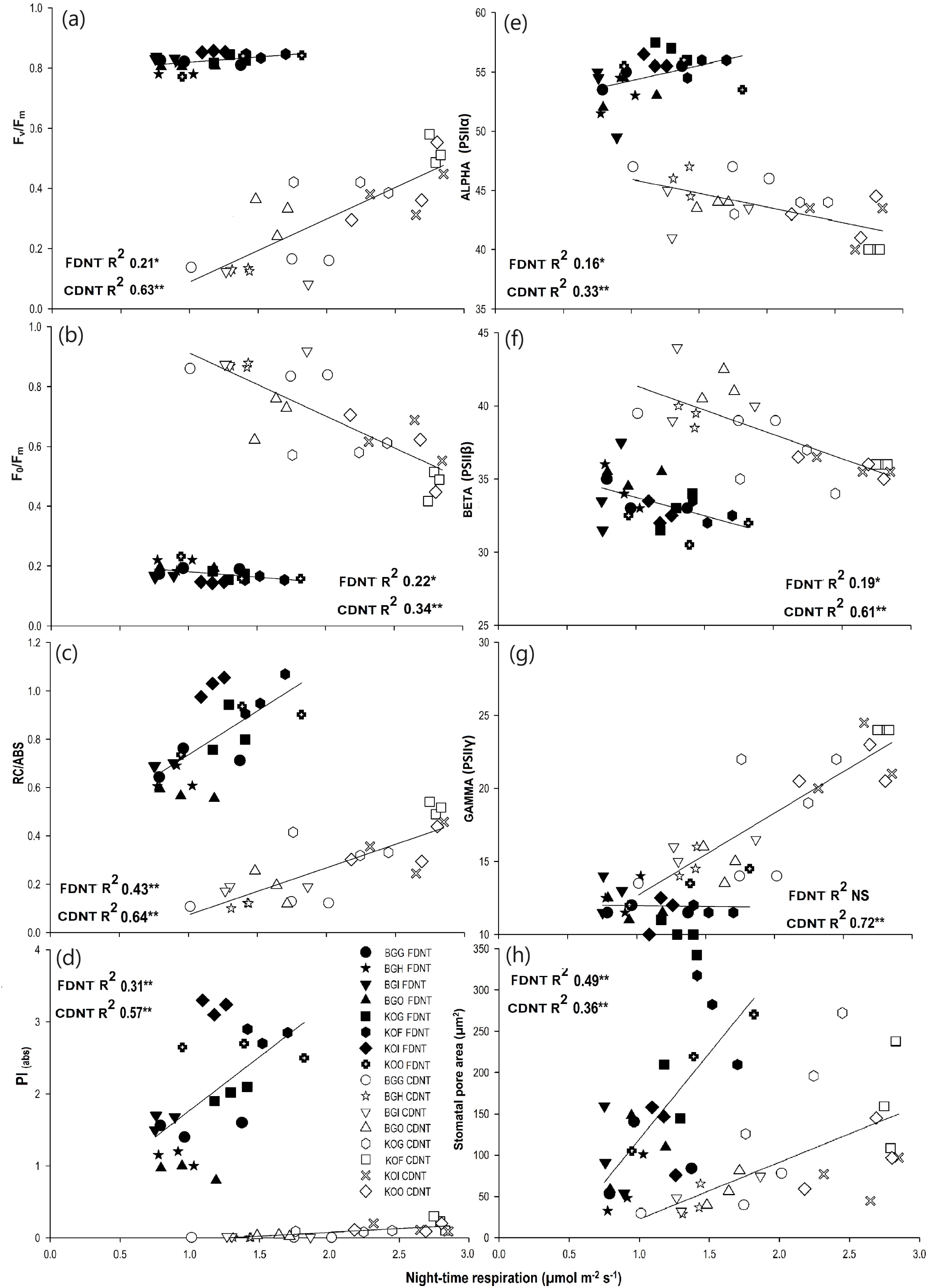
Correlation of night-time respiration with (A) F_v_/F_m_, (B) F_0_/F_m_, (C) RC/ABS, (D) PI_(abs)_ and antenna-size heterogeneity of PSII [(E) alpha (PSIIα), (F) beta (PSIIβ) and (G) gamma (PSIIγ)] and (H) stomatal pore area of different populations of *Bruguiera gymnorhiza* (BG) and *Kandelia obovata* (KO) under favourable (FDNT; *closed symbols*) and cold (CDNT [cold wave]; *open symbols*) day and night temperature conditions. Coefficient of determination (R^2^) followed by NS, * and ** corresponding to non-significance, significance at *P*<0.05 and *P*<0.01, respectively. BGG: BG from Guangxi, China, BGH: BG from Hainan, China, BGI: BG from Iriomote, Japan, BGO: BG from Okinawa, Japan, KOG: KO from Guangxi, China, KOF: KO from Fujian, China, KOI: KO from Iriomote, Japan and KOO: KO from Okinawa, Japan.

Under FDNT and CDNT, regardless of species and population, we found significant relationships. Specifically, there was a positive correlation of RC/ABS and PI _(abs)_ with F_v_/F_m_ and F_v_/F_0_ and a negative correlation with F_m_/F_0_ (Fig. 5). We also observed a positive correlation of absorptance with PSIIγ center, chlorophyll index, F_v_/F_m_, F_v_/F_0_, RC/ABS and PI _(abs)_ and a negative correlation with PSIIα and PSIIβ centers and F_0_/F_m_, while we found an opposite trend among transmittance and the above traits (Figs 6 and 7). We also found a negative correlation of the percentage of Q_B_ non-reducing centers with RC/ABS and F_v_/F_0_ (Fig. 8) and a positive correlation of stomatal pore area with F_v_/F_m_, F_v_/F_0_, RC/ABS and PI _(abs)_ (Fig. 9). Finally, we observed a positive correlation of night respiration with F_v_/F_m_, RC/ABS, PI_(abs),_ and stomatal pore area, and a negative correlation with F_0_/F_m_ and PSIIβ centers (Fig. 10).

## Discussion

Our results demonstrate that variation in the magnitude of tolerance of two mangrove species to a cold wave reflects an adaptation, which is evident from the diverse responses in photochemistry (light-energy utilization [chlorophyll fluorescence transients and P700 redox state]), stomatal functioning, PSII heterogeneity, and night respiration and variation in activation of important photoprotection mechanisms (CEF and NPQ). Meanwhile, we suggest that local adaptation takes part in determining the range of tolerance of the diverse populations from different latitudes to efficiently respond to the cold wave.

### Diversity in sensitivity of photochemistry and activation of photoprotection mechanisms during the cold wave

Significant changes in chlorophyll (Chl *a*) fluorescence transients (F_v_/F_m_, F_0_/F_m_, F_v_/F_0_, RC/ABS, Y[NPQ] and PI_(abs)_) and little change in the redox status of P700 of PSI (P_m_; oxidation of P700) in response to CDNT across populations of both mangrove species suggests the negative impact of a cold wave was greater on PSII than on PSI for both species (Table 3). The changes in Chl *a* fluorescence transients are interrelated, and during CDNT, these changes suggest reduced photochemical efficiency of PSII (F_v_/F_m_; Sunoj et al., 2016), damage of the thylakoid membrane (F_0_/F_m_; Sunoj et al., 2017) and water-splitting complex (F_v_/F_0_; Mathur et al., 2011a), failure of the photoprotection mechanisms of PSII (NPQ; Miyake, 2010), reduced electron transport between PSII and PSI (RC/ABS; Maswada et al., 2020), and decreased photosynthetic performance (PI_(abs)_; Strasser et al., 2004) which indicate the photoinhibition or photodamage of PSII (Table 3; Supplementary Table 2). However, the extent of the changes in photochemistry was much less in cold-tolerant *K. obovata* populations, particularly KOF from the coldest site, than in cold-susceptible *B. gymnorhiza* populations. Such distinctions in the interspecies sensitivity to cold are consistent with the observations of Chen et al., 2017 and Yan et al., 2021. Differences in the photochemistry of *K. obovata* and *B. gymnorhiza* in response to CDNT were further supported by the positive correlations of F_v_/F_m_ and F_v_/F_0_ and the negative correlation of F_0_/F_m_ with RC/ABS and PI _(abs)_ (Fig. 5). However, the magnitude of correlation between these traits varied under FDNT and CDNT (Fig. 5 and Table 3).

The sensitivity of PSII of *B. gymnorhiza* to a cold wave was further evident from the changes in the OJIP curve and energy pipeline leaf model (Figs 2 and 3). A distinct decline in the J-P phase of the Chl*a* transient curve indicates greater photoinhibition of PSII and decreased water-splitting, oxygen evolution, and electron transport through the plastoquinone pool (Mathur et al., 2021) was greater in *B. gymnorhiza* populations (Fig. 2). Similar to the OJIP curve, the energy pipeline leaf model provided evidence for greater negative impacts of CDNT on light absorptance and electron transport (decreased size of the arrow [ABS/CSo and ETo/CSo]), reaction centers (greater number of filled circles [TRo/CSo]) and energy dissipation (increased size of arrow [DIo/CSo]) in *B. gymnorhiza* populations (Mathur & Jajoo, 2014) (Fig. 3).

Photoinhibition of PSII during CDNT activated and influenced the rate of important photoprotection mechanisms across species. In our study, we observed an exceptional increase in CEF in the cold-tolerant population KOF from the colder site (Table 3). Previous studies have reported that cyclic electron flow (CEF) plays an important role in photoprotection during cold (Huang et al., 2010). The activation of CEF is a consequence of the photoinhibition of PSII, reduced electron transport between the photosystems, and limited photosynthesis (Agrawal et al., 2016; Neto et al., 2017). In KOF, activation of CEF was associated with an increase in NPQ (xanthophyll cycle; Muller et al., 2001; Miyake, 2010), which protected PSII and contributed to the cold tolerance of KOF as evident from the smaller decrease of F_v_/F_m_ and PI_(abs)_, respectively. Activation of NPQ is an important photoprotection mechanism bypassing the absorbed energy on PSII via heat dissipation to avoid overexcitation of PS II (Muller et al., 2001; Derks et al., 2015). Inactive or lower CEF in other populations could explain their low NPQ during the cold wave (Table 3).

### Reorganization of PSII heterogeneity for efficient responses to incident light, electron transport, and utilization of energy during a cold wave

The responses to incident light are very important under stress conditions because they can be beyond the capability of the plants to efficiently utilize the energy for photochemistry (Saez et al., 2019; Muller et al., 2001). Efficient responses to incident light alter the leaf optical properties (absorptance, reflectance, and transmittance), which slow down electron transport between PSII and PSI, and support the utilization of energy for CO_2_ fixation thus allowing photoprotection under cold (Miyake, 2020; Mathur et al., 2021; Sunoj et al., 2022).

We observed distinct variations and correlations in the leaf optical properties and PSII heterogeneity of both species (Table 3 and Fig 4 and 6). We observed less decrease in light absorptance, increase in light reflectance, and lower transmittance in *K. obovata* than in *B. gymnorhiza* during CDNT (Table 3 and Supplementary Table 3). Such changes in leaf optical properties can be due to the PSII antenna-size and reducing-side heterogeneity, chlorophyll degradation, and accumulation of epicuticular waxes (Laxman et al., 2013; Belgio et al., 2014; Marchin et al., 2014; Bukhanov et al., 2020).

PSII antenna-size heterogeneity is associated with three types of centers, namely, the alpha (α), beta (β) and gamma (γ) reaction centers (Hsu et al., 1989), classified according to their differences in PSII antenna size (number of chlorophyll molecules per PSII reaction centers). The PSIIα centers located in the grana partition regions are dominant, active, with larger antenna size, and responsible for most of the water-splitting activity and plastoquinone (PQ) reduction (electron transfer between the PSII reaction centers). The PSIIβ and PSIIγ centers located in the non-appressed region of the thylakoid membranes are less active, have smaller antenna sizes, and lack connectivity (energy transfer) (Anderson & Melis, 1983; Mathur et al., 2021). Under abiotic stress conditions, a reversible decrease in the number of active PSIIα centers and an increase in the number of less active PSIIβ and PSIIγ centers have been observed (Tomar et al., 2012; Mathur et al., 2021). Our study on sugarcane reported the reorganization of PSII active centers to inactive centers and the migration of PSIIβ and PSIIγ centers in a cold-tolerant genotype under cold (Mathur et al., 2021). We observed the same trend in PSII antenna-size heterogeneity in both *K. obovata* and *B. gymnorhiza* during CDNT. In *K. obovata*, the increase of PSIIγ centers was greater than that of PSIIβ centers and KOF from the colder site showed a greater percent increase in PSIIγ centers (Table 3, Supplementary Table 3 and Fig. 4).

The PSIIγ centers are smaller in size and less active than the other centers, possess the smallest antenna size (containing ∼ 50 Chl molecules and lacking inner LHC and peripheral LHC), and have a low trapping efficiency (Hsu & Lee, 1991,1995; Mathur et al., 2011a; Marchin et al., 2014). Hence, such an increase in PSIIγ centers during a cold wave can be attributed to allowing *K. obovata* to dissipate excess incident light by modulating the leaf optical properties and allowing only a small amount of absorbed energy to reach the photosystem reaction centers with the major portion of excess incident light being reflected (Table 3). This reduces the energy overload on the photosystems thus reducing photoinhibition of PSII and slowing down electron transport between photosystems to support the maintenance of oxidation of P700 in PSI (Sunoj et al., 2022) which is evident from the small changes in P_m_. This was further supported by the negative correlation of absorptance with active PSIIα and inactive PSIIβ centers, and its positive correlation with PSIIγ centers during CDNT with the opposite trend being observed for transmittance and antenna-size reaction centers (Fig. 6). The important role of PSIIγ centers in the efficient response, lower electron transport, utilization of light and photoprotection during CDNT which is further evident from positive correlation of absorptance with F_v_/F_m_, F_v_/F_0_, RC/ABS, and PI_(abs)_ (Fig. 7).

The optical properties of leaves also alter with existing biochemical composition, the concentration of light-harvesting pigments, water, and cell-wall structural materials in leaves (Ustin & Jacquemoud, 2020). A slight decrease in chlorophyll index (Table 3 and Supplementary Table 3) and correlations with absorptance (positive) and transmittance (negative) (Fig. 6D) during CDNT indicate the initial stage of chlorophyll degradation across species and populations. This implies the involvement of chlorophyll in the response to light by modulating the leaf optical properties along with PSII antenna-size heterogeneity (Carter & Knapp, 2001; Bauerle et al., 2004; Tang et al., 2015;; Mathur et al., 2021).

During CDNT, limiting electron transport between the photosystems was reinforced by the reorganization of the PSII reducing-side heterogeneity and it comprised the Q_B_-reducing and Q_B_-non-reducing centers. Q_B_-non-reducing centers are photochemically efficient but cannot transfer electrons efficiently from electron acceptor Q_A_− to secondary electron acceptor Q_B_ or reduce the PQ pool (Anderson & Melis, 1983; Mathur et al., 2011). The cold-tolerant *K. obovata* showed less reduction of active Q_B_-reducing centers (Table 3 and Supplementary Table 3) which can be attributed to the lower migration rate of active Q_B_-reducing centers from appressed to non-appressed parts of the thylakoid membrane. The migration of active Q_B_-reducing centers tends to increase the number of inactive Q_B_-nonreducing centers. The rate of electron transport from Q_A_ to Q_B_ is slower in Q_B_-nonreducing centers than active Q_B_-reducing centers. The increase in Q_B_-non-reducing centers diverts a major proportion of energy towards fluorescence and heat dissipation rather than photochemistry (Andersson & Melis, 1983; Mathur & Jajoo, 2020; Zhu et al., 2005).

Transformation of the Q_B_-reducing centers to Q_B_-non-reducing centers is often found under environmental stress conditions; this conversion is usually reversible, and the active PSII centers will recover to normal levels once the stress is relieved. The reversible deactivation of activity on the reducing side of PSII plays a photoprotective role in photosynthesis under environmental stresses (Strasser et al., 2004; Mathur et al., 2021). The positive correlation of RC/ABS with F_v_/F_m_ and F_v_/F_0_ and the negative correlation with F_0_/F_m_ (Fig. 4) and negative correlation of Q_B_-non-reducing centers with F_v_/F_0_ and RC/ABS (Fig. 8) suggest an important role of PSII reducing-side heterogeneity on the photochemical efficiency of PSII, reduction in electron release from water molecule to PSII, electron transport between the photosystems, and thylakoid membrane integrity during a cold wave (Supplementary Table 6).

### Contribution of stomatal functioning and role of night respiration to cope with a cold wave

Our study demonstrated that, in addition to the reorganization of PSII heterogeneity, stomatal functioning is distinctly varied due to a cold wave which is evident from the decrease in stomatal pore area of both species. The range of percentage reduction in the stomatal area during the cold wave was on par among the populations of *K. obovata* (26 to 50%) and *B. gymnorhiza* (28 to 47%) and the reduction was relatively low for KOF. However, the actual stomatal pore area was large in cold-tolerant *K. obovata* under FDNT and CDNT (Supplementary Table 4). Jurczyk et al. (2019) reported that cold-tolerant plant species tend to keep their stomata more open than sensitive species and thus maintain certain levels of photosynthesis, Rubisco activity, and NPQ under cold. However, the positive correlation of stomatal pore area with F_v_/F_m_, F_v_/F_0_, RC/ABS, PI_(abs)_ and night respiration, and the negative correlation with F_v_/F_0_ (Fig. 9) supports a role of stomatal functioning in coping with cold (Allen et al., 2000; Allen & Ort, 2001). Lower leaf temperature in *K. obovata* can be attributed to high stomatal pore area and transpiration which can avoid xylem failure and maintain certain level of photosynthesis (Roth-Nebelsick, 2007; Raven, 2014; Reef & Lovelock, 2015). Further studies are required to draw firm conclusions on the role of stomatal functioning.

A large increase in night respiration was found across mangrove species during CDNT with a difference in the magnitude (Table 3 and Supplementary Table 2). Night respiration is involved in the repair of damage to plants as a result of oxidative stress under different abiotic stresses (moderate to severe injuries) (Sunoj et al., 2016, 2020; Impa et al., 2018). The positive correlations of night respiration with F_v_/F_m_, RC/ABS and PI_(abs)_ and a negative correlation with F_0_/F_m_ validate the important role of night respiration in maintaining the integrity of thylakoid membranes, maintaining photochemistry (Fig. 10) and it can be attributed to the smaller CDNT impact on the photosystems of the tolerant *K. obovata*. Faster night respiration in *K. obovata* in our results is on par with Yan et al. (2021). Furthermore, the positive correlation of night respiration with other traits measured in this study viz., stomatal pore area and PSIIγ centers (Fig. 10) suggests that night respiration takes part in sustaining light-energy use and thereby contributes to photoprotection and photosynthetic integrity during cold waves.

The overall results imply that a cold wave negatively affected the photochemistry of mangroves and the impact was greater on PSII which activated photoprotection mechanisms (CEF and NPQ). The tolerance of mangroves to cold waves is genetically determined as evidenced by the distinct and diverse responses of tolerant and susceptible mangrove species. Reorganization of PSII heterogeneity and stomatal functioning along with faster night respiration allowed sustaining photochemistry and photoprotection by managing incident light, electron transport, and efficient utilization of energy and photosynthetic integrity during cold waves which enables the cold-tolerant *K. obovata* to be more resilient to cold waves. The minor impact of cold wave photochemistry and activation of important photoprotection mechanisms in KOF, the population from the coldest site (Fujian, China), can be attributed to the advantage of local adaptation to cold conditions in their natural habitats. This is the first study focused on PSII heterogeneity and stomatal functioning during a cold wave with an emphasis on its influence on light responses and energy utilization which extends our understanding of the adaption of mangrove species and their populations to cold waves.

## Acknowledgment

This work was supported by a Post-Doctoral Fellowship from Guangxi University, China granted to VSJS, by a visiting scholarship of Guangxi Science and Technology Department (GX2019007) and Post-Doctoral Fellowship for Women (PDFWM-2014-15-GEMAD-23945) from University Grants Commission (UGC), India to SM, and by a grant (31670406) of the National Natural Science Foundation of China and a Bagui Scholarship (C33600992001) granted to KFC.

## Author’s Contribution

The study was designed by KFC, VSJS, AJ, and SM. Experiments were performed by VSJS, SM, AWS, NIE, LY and ANAA. Analysis of the data and manuscript was drafted by VSJS and SM. The manuscript was edited and suggested critical comments by HL, KFC and AKSW. TK and AKSW collected and provided plant materials. VSJS and SM have contributed equally to the manuscript.

## Data availability

The data that support the findings of this study are openly available in figshare at https://doi.org/10.6084/m9.figshare.25333345.v2

## Abbreviations

CEF: cyclic electron flow
Fv/Fm: maximum quantum yield of PSII in the dark-adjusted state
Fv/F0: status of water splitting complex
F0/Fm: thylakoid membrane damage
PI(abs): performance index
Pm: maximum photo-oxidizable P700
RC/ABS: active PSII reaction centers per chlorophyll
Y(NPQ): effective quantum yield of non-photochemical quenching (NPQ).

---

**Supplementary Table 1:** List of various Chl*a* fluorescence and P700 transients.

| Parameters | Calculation | Description |
| --- | --- | --- |
| O | Fluorescence intensity at 50 μs | Fluorescence intensity when all RCs are open |
| J | Fluorescence intensity at 2 ms |  |
| I | Fluorescence intensity at 30 ms |  |
| P | Maximal fluorescence intensity |  |
| F<sub>m</sub> | Maximal fluorescence | Fluorescence intensity when all RCs are closed |
| F<sub>v</sub> | Maximal variable fluorescence, (F<sub>m</sub> – F<sub>50 μs</sub>) |  |
| RC/ABs | RC/TR<sub>o</sub>*TR<sub>o</sub>/ABS | The density of reaction center per chlorophyll |
| $\square_{Po/1} - \square_{Po}$ | F<sub>v</sub>/F<sub>0</sub> | Proportional to the activity of the water-splitting complex on the donor side of the PSII |
| $\psi_o$ | ET<sub>o</sub>/TR<sub>o</sub> | Probability that a trapped exciton moves an electron further than Q<sub>A</sub>– |
| PI<sub>(abs)</sub> | [RC/ABS] [$\square_{Po}/(1 - \square_{Po})$] [$\psi_o/(1 - \psi_o)$] | Performance Index on absorption basis |
| ABS/CS<sub>o</sub> approximated by F<sub>0</sub> |  | Absorption per cross section (CS) of the leaf tissue |
| TR<sub>o</sub>/CS<sub>o</sub> | (TR<sub>o</sub>/ABS)/(ABS/CS) | Trapping at time zero, per CS |
| ET<sub>o</sub>/CS<sub>o</sub> | (ET<sub>o</sub>/RC) (RC/CS) | Electron transport at time zero, per CS |
| D<sub>Io</sub>/C<sub>So</sub> | $(AB-/CS) - (TRo/CS)$ | Dissipation at time zero, per CS |
| Y(NPQ) | $1 - Y(II) - Y(NO)$ [$Y(II) = (F_m' - F_s)/F_m'$ and $Y(NO) = F_s/F_m'$] | Fraction of energy dissipated as heat through regulated non-photochemical quenching |
| CEF | ETRI-ETRII | Cyclic electron flow in PSI |
| P<sub>m</sub> |  | Maximum photo-oxidizable P700 |

**Supplementary Table 2:** The mean values (± SE) of chlorophyll *a* (Chl*a*) fluorescence transients, redox state of P700, cyclic electron flow and night respiration of the populations of two mangrove species (*Bruguiera gymnorhiza* [BG] and *Kandelia obovata* [KO]) under optimal (FDNT) and cold (CDNT; cold wave) day and night temperature conditions. BGG (BG from Guangxi, China), BGG: BG from Guangxi, China, BGH: BG from Hainan, China, BGI: BG from Iriomote, Japan, BGO: BG from Okinawa, Japan, KOG: KO from Guangxi, China, KOF: KO from Fujian, China, KOI: KO from Iriomote, Japan and KOO: KO from Okinawa, Japan. F_v_/F_m_: maximum quantum yield of PSII in the dark-adjusted state (relative units); F_v_/F_0_: status of water splitting complex (relative units), F_0_/F_m_: thylakoid membrane damage (relative units); RC/ABS: active PSII reaction centers per chlorophyll; PI_(abs)_:performance index; Y(NPQ): effective quantum yield of NPQ; P_m_: maximum photo-oxidizable P700 (ΔI/I); CEF: cyclic electron flow.

| Species/ Populations/<br>Temperature conditions | $F_v/F_m$ | $F_v/F_0$ | $F_0/F_m$ | RC/ABS | $PI_{(abs)}$ | Y(NPQ) | $P_m$ | CEF | Night<br>respiration<br>($\mu\text{mol m}^{-2}$<br>$\text{s}^{-1}$) |
| --- | --- | --- | --- | --- | --- | --- | --- | --- | --- |
| *Bruguiera gymnorhiza* (BG) |  |  |  |  |  |  |  |  |  |
| FDNT |  |  |  |  |  |  |  |  |  |
| BGG | 0.82<br>$\pm 0.005^{\text{a}\square}$ | 4.41<br>$\pm 0.18^{\text{ab}}$ | 0.19<br>$\pm 0.01^{\text{ab}}$ | 0.71<br>$\pm 0.03^{\text{a}}$ | 1.56<br>$\pm 0.02^{\text{a}}$ | 0.44<br>$\pm 0.04^{\text{b}}$ | 3.7<br>$\pm 0.1^{\text{ab}}$ | 13.53<br>$\pm 1.98^{\text{bc}}$ | 1.04<br>$\pm 0.17^{\text{a}}$ |
| BGH | 0.79<br>$\pm 0.012^{\text{a}}$ | 3.82<br>$\pm 0.31^{\text{c}}$ | 0.21<br>$\pm 0.01^{\text{a}}$ | 0.64<br>$\pm 0.03^{\text{b}}$ | 1.15<br>$\pm 0.02^{\text{ab}}$ | 0.47<br>$\pm 0.01^{\text{ab}}$ | 3.4<br>$\pm 0.2^{\text{ab}}$ | 12.14<br>$\pm 1.29^{\text{c}}$ | 0.91<br>$\pm 0.07^{\text{a}}$ |
| BGI | 0.83<br>$\pm 0.002^{\text{a}}$ | 5.04<br>$\pm 0.08^{\text{a}}$ | 0.17<br>$\pm 0.001^{\text{bc}}$ | 0.69<br>$\pm 0.001^{\text{a}}$ | 1.68<br>$\pm 0.02^{\text{a}}$ | 0.50<br>$\pm 0.03^{\text{a}}$ | 4.1<br>$\pm 0.1^{\text{a}}$ | 14.25<br>$\pm 1.02^{\text{b}}$ | 0.80<br>$\pm 0.05^{\text{b}}$ |
| BGO | 0.81<br>$\pm 0.001^{\text{a}}$ | 4.16<br>$\pm 0.05^{\text{ab}}$ | 0.19<br>$\pm 0.05^{\text{ab}}$ | 0.57<br>$\pm 0.01^{\text{c}}$ | 0.97<br>$\pm 0.01^{\text{b}}$ | 0.38<br>$\pm 0.03^{\text{c}}$ | 4.0<br>$\pm 0.4^{\text{a}}$ | 18.40<br>$\pm 2.02^{\text{a}}$ | 0.97<br>$\pm 0.07^{\text{a}}$ |
| CDNT |  |  |  |  |  |  |  |  |  |
| BGG | 0.15<br>$\pm 0.009^{\text{c}}$ | 0.18<br>$\pm 0.01^{\text{b}}$ | 0.84<br>$\pm 0.01^{\text{a}}$ | 0.12<br>$\pm 0.01^{\text{c}}$ | 0.006<br>$\pm 0.001^{\text{c}}$ | 0.33<br>$\pm 0.05^{\text{a}}$ | 3.6<br>$\pm 0.3^{\text{a}}$ | 13.51<br>$\pm 2.57^{\text{a}}$ | 1.59<br>$\pm 0.30^{\text{ab}}$ |
| BGH | 0.13<br>$\pm 0.003^{\text{c}}$ | 0.15<br>$\pm 0.001^{\text{c}}$ | 0.87<br>$\pm 0.001^{\text{a}}$ | 0.11<br>$\pm 0.01^{\text{c}}$ | 0.005<br>$\pm 0.001^{\text{c}}$ | 0.26<br>$\pm 0.09^{\text{b}}$ | 3.4<br>$\pm 0.4^{\text{a}}$ | 11.07<br>$\pm 1.09^{\text{b}}$ | 1.39<br>$\pm 0.04^{\text{b}}$ |
| BGI | 0.27 | 0.38 | 0.73 | 0.18 | 0.016 | 0.33 | 3.3 | 9.80 | 1.48 |
|  | ±0.015<sup>b</sup> | ±0.04<sup>a</sup> | ±0.02<sup>b</sup> | ±0.01<sup>b</sup> | ±0.001<sup>b</sup> | ±0.04<sup>a</sup> | ±0.4<sup>ab</sup> | ±2.1<sup>c</sup> | ±0.19<sup>ab</sup> |
| BGO | 0.33<br>±0.04<sup>a</sup> | 0.36<br>±0.05<sup>a</sup> | 0.66<br>±0.04<sup>c</sup> | 0.022<br>±0.04<sup>a</sup> | 0.032<br>±0.001<sup>a</sup> | 0.18<br>±0.05<sup>c</sup> | 3.7<br>±0.2<sup>a</sup> | 13.07<br>±0.38<sup>a</sup> | 1.61<br>±0.07<sup>a</sup> |
| *Kandelia obovata* (KO) |  |  |  |  |  |  |  |  |  |
| FDNT |  |  |  |  |  |  |  |  |  |
| KOG | 0.83<br>±0.008<sup>a</sup> | 5.35<br>±0.30<sup>a</sup> | 0.17<br>±0.01<sup>a</sup> | 0.83<br>±0.06<sup>ab</sup> | 2.02<br>±0.3<sup>bc</sup> | 0.53<br>±0.02<sup>a</sup> | 4.2<br>±0.2<sup>a</sup> | 15.07<br>±1.83<sup>b</sup> | 1.29<br>±0.02<sup>a</sup> |
| KOF | 0.84<br>±0.004<sup>a</sup> | 4.85<br>±0.16<sup>ab</sup> | 0.15<br>±0.001<sup>a</sup> | 0.97<br>±0.05<sup>a</sup> | 2.8<br>±0.2<sup>ab</sup> | 0.36<br>±0.07<sup>bc</sup> | 4.6<br>±0.3<sup>a</sup> | 11.08<br>±4.41<sup>c</sup> | 1.55<br>±0.05<sup>a</sup> |
| KOI | 0.85<br>±0.001<sup>a</sup> | 5.88<br>±0.06<sup>a</sup> | 0.14<br>±0.001<sup>a</sup> | 1.02<br>±0.02<sup>a</sup> | 3.24<br>±0.02<sup>a</sup> | 0.44<br>±0.01<sup>ab</sup> | 3.9<br>±0.1<sup>ab</sup> | 16.52<br>±1.19<sup>b</sup> | 1.18<br>±0.25<sup>a</sup> |
| KOO | 0.84<br>±0.023<sup>a</sup> | 4.81<br>±0.54<sup>ab</sup> | 0.15<br>±0.02<sup>a</sup> | 0.94<br>±0.06<sup>a</sup> | 2.65<br>±0.3<sup>ab</sup> | 0.45<br>±0.01<sup>ab</sup> | 3.6<br>±0.1<sup>ab</sup> | 23.38<br>±0.14<sup>a</sup> | 1.38<br>±0.08<sup>a</sup> |
| CDNT |  |  |  |  |  |  |  |  |  |
| KOG | 0.42<br>±0.012<sup>b</sup> | 1.01<br>±0.03<sup>a</sup> | 0.57<br>±0.01<sup>a</sup> | 0.36<br>±0.03<sup>c</sup> | 0.09<br>±0.002<sup>b</sup> | 0.31<br>±0.02<sup>a</sup> | 4.1<br>±0.2<sup>a</sup> | 13.99<br>±2.56<sup>b</sup> | 2.15<br>±0.02<sup>a</sup> |
| KOF | 0.53<br>±0.028<sup>a</sup> | 1.17<br>±0.13<sup>a</sup> | 0.46<br>±0.03<sup>b</sup> | 0.52<br>±0.01<sup>a</sup> | 0.23<br>±0.003<sup>a</sup> | 0.39<br>±0.07<sup>a</sup> | 4.4<br>±0.2<sup>a</sup> | 17.82<br>±3.99<sup>a</sup> | 2.79<br>±0.16<sup>a</sup> |
| KOI | 0.41<br>±0.039<sup>b</sup> | 0.76<br>±0.10<sup>b</sup> | 0.59<br>±0.04<sup>a</sup> | 0.40<br>±0.06<sup>b</sup> | 0.11<br>±0.003<sup>b</sup> | 0.14<br>±0.01<sup>b</sup> | 4.1<br>±0.4<sup>a</sup> | 13.20<br>±1.09<sup>b</sup> | 2.61<br>±0.19<sup>a</sup> |
| KOO | 0.45<br>±0.077<sup>b</sup> | 0.83<br>±0.26<sup>b</sup> | 0.55<br>±0.08<sup>a</sup> | 0.38<br>±0.05<sup>c</sup> | 0.11<br>±0.001<sup>b</sup> | 0.39<br>±0.01<sup>a</sup> | 3.2<br>±0.3<sup>b</sup> | 18.59<br>±2.82<sup>a</sup> | 2.56<br>±0.10<sup>a</sup> |
| Mean BG and KO |  |  |  |  |  |  |  |  |  |
| BG (FDNT) | 0.81<br>±0.01 | 4.36<br>±0.25 | 0.19<br>±0.01 | 0.65<br>±0.03 | 1.34<br>±0.17 | 0.45<br>±0.03 | 3.8<br>±0.16 | 14.6<br>±3.1 | 0.93<br>±0.05 |
| BG (CDNT) | 0.22 | 0.27 | 0.78 | 0.16 | 0.01 | 0.28 | 3.5 | 11.9 | 1.52 |
| | $\pm 0.05^{**}$ | $\pm 0.16^{**}$ | $\pm 0.04^{**}$ | $\pm 0.02^{**}$ | $\pm 0.01^{**}$ | $\pm 0.04^{**}$ | $\pm 0.09^*$ | $\pm 0.9^*$ | $\pm 0.05^{**}$ |
| BG percentage (%) change | **-73** | **-94** | **311** | **-75** | **-99** | **-38** | **-8** | **-18** | **63** |
| KO (FDNT) | 0.84<sup>ns</sup><br>$\pm 0.01$ | 5.22<sup>A</sup><br>$\pm 0.24$ | 0.15<sup>A</sup><br>$\pm 0.01$ | 0.94<sup>AA</sup><br>$\pm 0.04$ | 2.7<sup>AA</sup><br>$\pm 0.25$ | 0.43<sup>ns</sup><br>$\pm 0.04$ | 4.25<sup>ns</sup><br>$\pm 0.3$ | 16.5<sup>ns</sup><br>$\pm 2.6$ | 1.35<sup>A</sup><br>$\pm 0.08$ |
| KO (CDNT) | 0.45<sup>AA</sup><br>$\pm 0.03^{**}$ | 0.94<sup>AA</sup><br>$\pm 0.12^{**}$ | 0.54<sup>AA</sup><br>$\pm 0.03^{**}$ | 0.42<sup>AA</sup><br>$\pm 0.04^{**}$ | 0.14<sup>AA</sup><br>$\pm 0.03^{**}$ | 0.32<sup>A</sup><br>$\pm 0.07^{**}$ | 4.0<sup>A</sup><br>$\pm 0.3^{**}$ | 15.9<sup>A</sup><br>$\pm 1.3^*$ | 2.53<sup>AA</sup><br>$\pm 0.1^{**}$ |
| KO percentage (%) change | **-46** | **-82** | **260** | **-55** | **-95** | **-26** | **-7** | **-4** | **87** |

Favorable day and night temperatures (FDNT); cold day and night temperatures (CDNT [cold wave]); mean values of different populations re followed by different lowercase letters are significantly different according to Duncan's multiple range test (DMRT; $P < 0.01$ ; the test was conducted independently for individual species under each temperature condition). The mean values of individual species (irrespective of four populations) under different temperature conditions and mean values of different species (BG and KO; mean value of all four populations) under similar temperature conditions were compared using student's t-test and \* and <sup>A</sup> and \*\* and <sup>AA</sup> corresponding to significance at $P < 0.05$ and $P < 0.01$ , respectively. Percentage (%) change indicates the difference differences from FDNT to CDNT. □ Values are ± standard error of respective mean values. ns denotes non-significant.

**Supplementary Table 3:** The mean values (± SE) of PSII heterogeneity, leaf optical properties and chlorophyll index of the populations of two mangrove species (*Bruguiera gymnorhiza* [BG] and *Kandelia obovata* [KO]) under favorable (FDNT) and cold (CDNT; cold wave) day and night temperature conditions. BGG: BG from Guangxi, China, BGH: BG from Hainan, China, BGI: BG from Iriomote, Japan, BGO: BG from Okinawa, Japan, KOG: KO from Guangxi, China, KOF: KO from Fujian, China, KOI: KO from Iriomote, Japan and KOO: KO from Okinawa, Japan.

| PSII heterogeneity |  |  |  |  |  | Reflectance (%) | Transmittance (%) | Absorptance (%) | Chlorophyll index (SPAD values) |
| --- | --- | --- | --- | --- | --- | --- | --- | --- | --- |
| Reducing-side heterogeneity |  |  | Antenna-size heterogeneity (%) |  |  |  |  |  |  |
| Species/<br>Populations/<br>Temperature<br>conditions | % of Q<sub>B</sub>-<br>reducing<br>Centers<br>(1-Bo) | % of Q<sub>B</sub>-<br>non-reducing<br>Centers (Bo) | $\alpha$ | $\beta$ | $\gamma$ | | | | |
| *Bruguiera gymnorhiza* (BG) |  |  |  |  |  |  |  |  |  |
| FDNT |  |  |  |  |  |  |  |  |  |
| BGG | 53<br>$\pm 0.8^{ab\Box}$ | 47<br>$\pm 0.6^a$ | 54<br>$\pm 0.4^a$ | 35<br>$\pm 0.7^{ab}$ | 11<br>$\pm 0.7^{ab}$ | 17<br>$\pm 1.0^a$ | 20<br>$\pm 1.5^a$ | 63<br>$\pm 2.1^b$ | 49<br>$\pm 1.0^{ab}$ |
| BGH | 54<br>$\pm 0.8^{ab}$ | 46<br>$\pm 0.4^a$ | 53<br>$\pm 0.6^a$ | 34<br>$\pm 0.7^{ab}$ | 13<br>$\pm 0.8^a$ | 13<br>$\pm 1.2^b$ | 22<br>$\pm 1.0^a$ | 65<br>$\pm 1.4^{ab}$ | 52<br>$\pm 0.8^a$ |
| BGI | 61<br>$\pm 0.8^a$ | 39<br>$\pm 0.5^b$ | 49<br>$\pm 1.1^{ab}$ | 37<br>$\pm 0.8^a$ | 14<br>$\pm 0.5^a$ | 15<br>$\pm 0.1^a$ | 18<br>$\pm 0.9^b$ | 67<br>$\pm 0.9^{ab}$ | 49<br>$\pm 1.0^{ab}$ |
| BGO | 60<br>$\pm 0.8^a$ | 40<br>$\pm 0.2^{ab}$ | 53<br>$\pm 1.7^a$ | 35<br>$\pm 1.4^{ab}$ | 12<br>$\pm 0.7^{ab}$ | 15<br>$\pm 1.2^a$ | 14<br>$\pm 1.4^c$ | 71<br>$\pm 0.8^a$ | 52<br>$\pm 1.5^a$ |
| CDNT |  |  |  |  |  |  |  |  |  |
| BGG | 34<br>$\pm 0.6^b$ | 66<br>$\pm 0.7^a$ | 47<br>$\pm 0.4^a$ | 39<br>$\pm 0.3^c$ | 14<br>$\pm 0.3^{ab}$ | 14<br>$\pm 0.6^a$ | 26<br>$\pm 1.0^a$ | 60<br>$\pm 1.0^b$ | 46<br>$\pm 0.8^c$ |
| BGH | 33 | 67 | 46 | 39 | 15 | 13 | 26 | 61 | 50 |
| | $\pm 0.6^b$ | $\pm 0.8^a$ | $\pm 0.8^a$ | $\pm 1.0^c$ | $\pm 0.8^a$ | $\pm 0.5^a$ | $\pm 0.5^a$ | $\pm 1.0^b$ | $\pm 0.5^a$ |
| BGI | 49<br>$\pm 0.7^a$ | 51<br>$\pm 0.9^b$ | 40<br>$\pm 0.6^b$ | 45<br>$\pm 0.6^a$ | 15<br>$\pm 0.5^a$ | 14<br>$\pm 1.0^a$ | 27<br>$\pm 0.7^a$ | 59<br>$\pm 0.7^b$ | 48<br>$\pm 1.3^b$ |
| BGO | 48<br>$\pm 0.6^b$ | 52<br>$\pm 0.4^b$ | 44<br>$\pm 1.0^{ab}$ | 40<br>$\pm 1.1^b$ | 16<br>$\pm 0.4^a$ | 10<br>$\pm 1.5^b$ | 25<br>$\pm 0.8^a$ | 65<br>$\pm 1.0^a$ | 51<br>$\pm 0.8^a$ |
| *Kandelia obovata* (KO) |  |  |  |  |  |  |  |  |  |
| FDNT |  |  |  |  |  |  |  |  |  |
| KOG | 60<br>$\pm 0.4^a$ | 40<br>$\pm 0.4^a$ | 56<br>$\pm 1.0^a$ | 32<br>$\pm 0.8^a$ | 12<br>$\pm 0.5^a$ | 15<br>$\pm 0.9^a$ | 13<br>$\pm 0.5^a$ | 72<br>$\pm 1.5^{ab}$ | 54<br>$\pm 0.7^a$ |
| KOF | 61<br>$\pm 0.7^a$ | 39<br>$\pm 0.3^a$ | 58<br>$\pm 0.4^a$ | 33<br>$\pm 0.3^a$ | 9<br>$\pm 0.3^b$ | 14<br>$\pm 1.0^a$ | 11<br>$\pm 0.1^a$ | 75<br>$\pm 2.1^a$ | 52<br>$\pm 1.2^{ab}$ |
| KOI | 64<br>$\pm 0.8^a$ | 36<br>$\pm 0.4^{ab}$ | 57<br>$\pm 0.7^a$ | 32<br>$\pm 0.3^a$ | 11<br>$\pm 0.7^a$ | 14<br>$\pm 1.1^a$ | 13<br>$\pm 1.4^a$ | 73<br>$\pm 0.7^{ab}$ | 50<br>$\pm 2.0^b$ |
| KOO | 61<br>$\pm 0.4^a$ | 39<br>$\pm 0.7^a$ | 55<br>$\pm 0.7^a$ | 32<br>$\pm 0.4^a$ | 13<br>$\pm 0.7^a$ | 12<br>$\pm 0.6^b$ | 13<br>$\pm 0.9^a$ | 75<br>$\pm 0.1^a$ | 54<br>$\pm 1.0^a$ |
| CDNT |  |  |  |  |  |  |  |  |  |
| KOG | 46<br>$\pm 0.3^{ab}$ | 54<br>$\pm 0.2^a$ | 40<br>$\pm 1.0^b$ | 36<br>$\pm 1.1^a$ | 24<br>$\pm 1.0^a$ | 16<br>$\pm 1.0^a$ | 14<br>$\pm 0.3^{ab}$ | 70<br>$\pm 1.6^b$ | 53<br>$\pm 0.9^a$ |
| KOF | 48<br>$\pm 0.6^a$ | 52<br>$\pm 0.6^b$ | 46<br>$\pm 0.3^a$ | 34<br>$\pm 0.4^a$ | 20<br>$\pm 0.2^a$ | 16<br>$\pm 1.0^a$ | 10<br>$\pm 1.2^c$ | 74<br>$\pm 1.0^a$ | 53<br>$\pm 1.4^a$ |
| KOI | 44<br>$\pm 0.7^b$ | 56<br>$\pm 0.5^a$ | 40<br>$\pm 0.8^b$ | 36<br>$\pm 0.3^a$ | 24<br>$\pm 0.9^a$ | 13<br>$\pm 1.0^b$ | 16<br>$\pm 0.9^a$ | 71<br>$\pm 0.7^b$ | 51<br>$\pm 1.0^b$ |
| KOO | 47<br>$\pm 0.5^a$ | 53<br>$\pm 0.9^b$ | 43<br>$\pm 0.7^a$ | 37<br>$\pm 0.4^a$ | 21<br>$\pm 0.6^a$ | 12<br>$\pm 0.8^b$ | 15<br>$\pm 0.4^a$ | 73<br>$\pm 1.0^a$ | 51<br>$\pm 0.1^b$ |
| Mean BG and KO |  |  |  |  |  |  |  |  |  |
| BG (FDNT) | 57 | 43 | 52 | 35 | 13 | 15 | 19 | 67 | 51 |
| | $\pm 2.3$ | $\pm 1.3$ | $\pm 0.5$ | $\pm 0.3$ | $\pm 0.5$ | $\pm 0.8$ | $\pm 1.7$ | $\pm 1.7$ | $\pm 0.9$ |
| BG (CDNT) | 41<br>$\pm 6.1^{**}$ | 59<br>$\pm 5.5^{**}$ | 44<br>$\pm 0.9^{**}$ | 41<br>$\pm 0.6^{**}$ | 15<br>$\pm 0.4^*$ | 13<br>$\pm 0.9^*$ | 26<br>$\pm 0.4^{**}$ | 61<br>$\pm 1.3^*$ | 49<br>$\pm 1.1^*$ |
| BG percentage (%)<br>change | **-28** | **37** | **-15** | **17** | **15** | **-13** | **37** | **-9** | **-4** |
| KO (FDNT) | 62<sup>A</sup><br>$\pm 0.9$ | 39<sup>A</sup><br>$\pm 0.9$ | 57<sup>A</sup><br>$\pm 0.4$ | 32<sup>ns</sup><br>$\pm 0.3$ | 11<sup>ns</sup><br>$\pm 0.6$ | 14<sup>ns</sup><br>$\pm 0.6$ | 13<sup>A</sup><br>$\pm 0.5$ | 74<sup>A</sup><br>$\pm 0.8$ | 53<sup>ns</sup><br>$\pm 1.0$ |
| KO (CDNT) | 46<sup>A</sup><br>$\pm 1.2^{**}$ | 54<sup>A</sup><br>$\pm 1.7^{**}$ | 42<sup>A</sup><br>$\pm 0.9^{**}$ | 36<sup>A</sup><br>$\pm 0.3^*$ | 22<sup>AA</sup><br>$\pm 0.7^{**}$ | 14<sup>ns</sup><br>$\pm 1.0^{ns}$ | 14<sup>AA</sup><br>$\pm 1.3^*$ | 72<sup>A</sup><br>$\pm 0.9^*$ | 52<sup>A</sup><br>$\pm 0.6^*$ |
| KO percentage (%)<br>change | **-26** | **38** | **-26** | **13** | **100** | **3** | **8** | **-3** | **-2** |

Favorable day and night temperatures (FDNT); cold day and night temperatures (CDNT; cold wave); mean values of populations followed by different lowercase letters are significantly different according to Duncan's multiple range test (DMRT; $P < 0.01$ ; the test was conducted independently for individual species under each temperature conditions). The mean values of individual species (irrespective of four populations) under different temperature conditions and mean values of different species (BG and KO; mean value of all four populations) under similar temperature conditions were compared using student's t-test and \* and <sup>A</sup> and \*\* and <sup>AA</sup> corresponding to significance at $P < 0.05$ and $P < 0.01$ , respectively. Percentage (%) change indicates the difference differences from FDNT to CDNT. □ Values are ± standard error of respective mean values. ns denotes non-significant.

**Supplementary Table 4:** The mean values (±SE) of stomatal aperture and leaf temperature of the populations of two mangrove species (*Bruguiera gymnorhiza* [BG] and *Kandelia obovata* [KO]) under favorable (FDNT) and cold (CDNT; cold wave) day and night temperature conditions. BGG: BG from Guangxi, China, BGH: BG from Hainan, China, BGI: BG from Iriomote, Japan, BGO: BG from Okinawa, Japan, KOG: KO from Guangxi, China, KOF: KO from Fujian, China, KOI: KO from Iriomote, Japan and KOO: KO from Okinawa, Japan.

| Species/ Populations/<br>Temperature conditions | Stomatal pore<br>length ($\mu\text{m}$) | Stomatal pore<br>width ($\mu\text{m}$) | Stomatal pore area<br>($\mu\text{m}^2$) | Leaf temperature<br>($^{\circ}\text{C}$) |
| --- | --- | --- | --- | --- |
| *Bruguiera gymnorhiza* (BG) |  |  |  |  |
| FDNT |  |  |  |  |
| BGG | 23.8<br>$\pm 0.16^{\text{b}\square}$ | 5.1<br>$\pm 0.15^{\text{a}}$ | 92.5<br>$\pm 25.4^{\text{a}}$ | 29.3<br>$\pm 0.22^{\text{a}}$ |
| BGH | 28.2<br>$\pm 0.19^{\text{a}}$ | 2.8<br>$\pm 0.12^{\text{b}}$ | 60.5<br>$\pm 2.4^{\text{b}}$ | 30.9<br>$\pm 0.12^{\text{a}}$ |
| BGI | 28.9<br>$\pm 0.19^{\text{a}}$ | 4.7<br>$\pm 0.15^{\text{a}}$ | 101.2<br>$\pm 3.2^{\text{a}}$ | 29.8<br>$\pm 0.20^{\text{a}}$ |
| BGO | 27.5<br>$\pm 0.22^{\text{a}}$ | 5.7<br>$\pm 0.14^{\text{a}}$ | 105.2<br>$\pm 2.7^{\text{a}}$ | 30.7<br>$\pm 0.20^{\text{a}}$ |
| CDNT |  |  |  |  |
| BGG | 23.2<br>$\pm 0.14^{\text{c}}$ | 2.7<br>$\pm 0.10^{\text{a}}$ | 49.0<br>$\pm 1.8^{\text{bc}}$ | 14.0<br>$\pm 0.21^{\text{a}}$ |
| BGH | 26.1<br>$\pm 0.22^{\text{ab}}$ | 1.9<br>$\pm 0.60^{\text{c}}$ | 43.4<br>$\pm 1.2^{\text{c}}$ | 12.5<br>$\pm 0.35^{\text{b}}$ |
| BGI | 27.2<br>$\pm 0.17^{\text{a}}$ | 2.3<br>$\pm 0.53^{\text{b}}$ | 51.6<br>$\pm 1.3^{\text{b}}$ | 15.0<br>$\pm 0.24^{\text{a}}$ |
| BGO | 25.0<br>$\pm 0.16^{\text{b}}$ | 2.9<br>$\pm 0.7^{\text{a}}$ | 58.8<br>$\pm 1.3^{\text{a}}$ | 12.3<br>$\pm 0.55^{\text{b}}$ |
| *Kandelia obovata* (KO) |  |  |  |  |
| FDNT |  |  |  |  |
| KOG | 31.6<br>±0.94<sup>a</sup> | 11.6±<br>2.38<sup>a</sup> | 266.7<br>±58.2<sup>a</sup> | 29.0<br>±0.42<sup>a</sup> |
| KOF | 31.9<br>±0.30<sup>a</sup> | 9.6<br>±1.16<sup>ab</sup> | 228.4<br>±31.8<sup>ab</sup> | 28.5<br>±0.32<sup>a</sup> |
| KOI | 31.8<br>±0.28<sup>a</sup> | 5.5<br>±0.24<sup>b</sup> | 129.5<br>±4.5<sup>c</sup> | 29.2<br>±0.32<sup>a</sup> |
| KOO | 33.1<br>±0.35<sup>a</sup> | 8.1<br>±0.25<sup>ab</sup> | 200.4<br>±5.9<sup>b</sup> | 29.5<br>±0.23<sup>a</sup> |
| CDNT |  |  |  |  |
| KOG | 31.3<br>±0.32<sup>a</sup> | 8.9<br>±0.2<sup>a</sup> | 198.6<br>±4.5<sup>a</sup> | 13.1<br>±0.49<sup>a</sup> |
| KOF | 31.0<br>±0.31<sup>a</sup> | 7.3<br>±0.2<sup>b</sup> | 169.3<br>±4.0<sup>b</sup> | 11.8<br>±0.18<sup>b</sup> |
| KOI | 29.6<br>±0.36<sup>ab</sup> | 2.9<br>±0.73<sup>c</sup> | 72.7<br>±2.5<sup>d</sup> | 13.9<br>±0.50<sup>a</sup> |
| KOO | 32.4<br>±0.34<sup>a</sup> | 3.9<br>±0.12<sup>c</sup> | 100.5<br>±2.7<sup>c</sup> | 13.8<br>±0.54<sup>a</sup> |
| Mean BG and KO |  |  |  |  |
| BG (FDNT) | 27.1<br>±1.1 | 4.6<br>±0.6 | 89.9<br>±10.1 | 30.2<br>±0.4 |
| BG(CDNT) | 25.4<br>±0.9* | 2.5<br>±0.2** | 50.7<br>±3.2** | 13.5<br>±0.6** |
| BG percentage (%) change | **-6** | **-46** | **-44** | **-55** |
| KO (FDNT) | 31.9<sup>A</sup><br>±0.1 | 9.6<sup>AA</sup><br>±1.3 | 206.3<sup>AA</sup><br>±29.0 | 29.1<sup>ns</sup><br>±0.2 |
| KO (CDNT) | 31.1<sup>A</sup><br>±0.6<sup>ns</sup> | 5.8<sup>AA</sup><br>±1.4<sup>**</sup> | 135.3<sup>AA</sup><br>±29.3<sup>**</sup> | 13.2<sup>ns</sup><br>±0.5<sup>**</sup> |
| KO percentage (%) change | **-3** | **-40** | **-34** | **-55** |

Favorable day and night temperatures (FDNT); cold day and night temperatures (CDNT; cold wave); Mean values of different populations followed by different lowercase letters are significantly different according to Duncan's multiple range test (DMRT; $P < 0.01$ ; the test was conducted independently for individual species under each temperature condition). The mean values of individual species (irrespective of four populations) under different temperature conditions and mean values of different species (BG and KO; mean value of all four populations) under each temperature conditions were compared using student's t-test and \* and <sup>A</sup> and \*\* and <sup>AA</sup> corresponding to significance at $P < 0.05$ and $P < 0.01$ , respectively. Percentage (%) change indicate the difference differences from FDNT to CDNT. □ Values are ± standard error of respective mean values. ns denotes non-significant.

**Supplementary Table 5:**
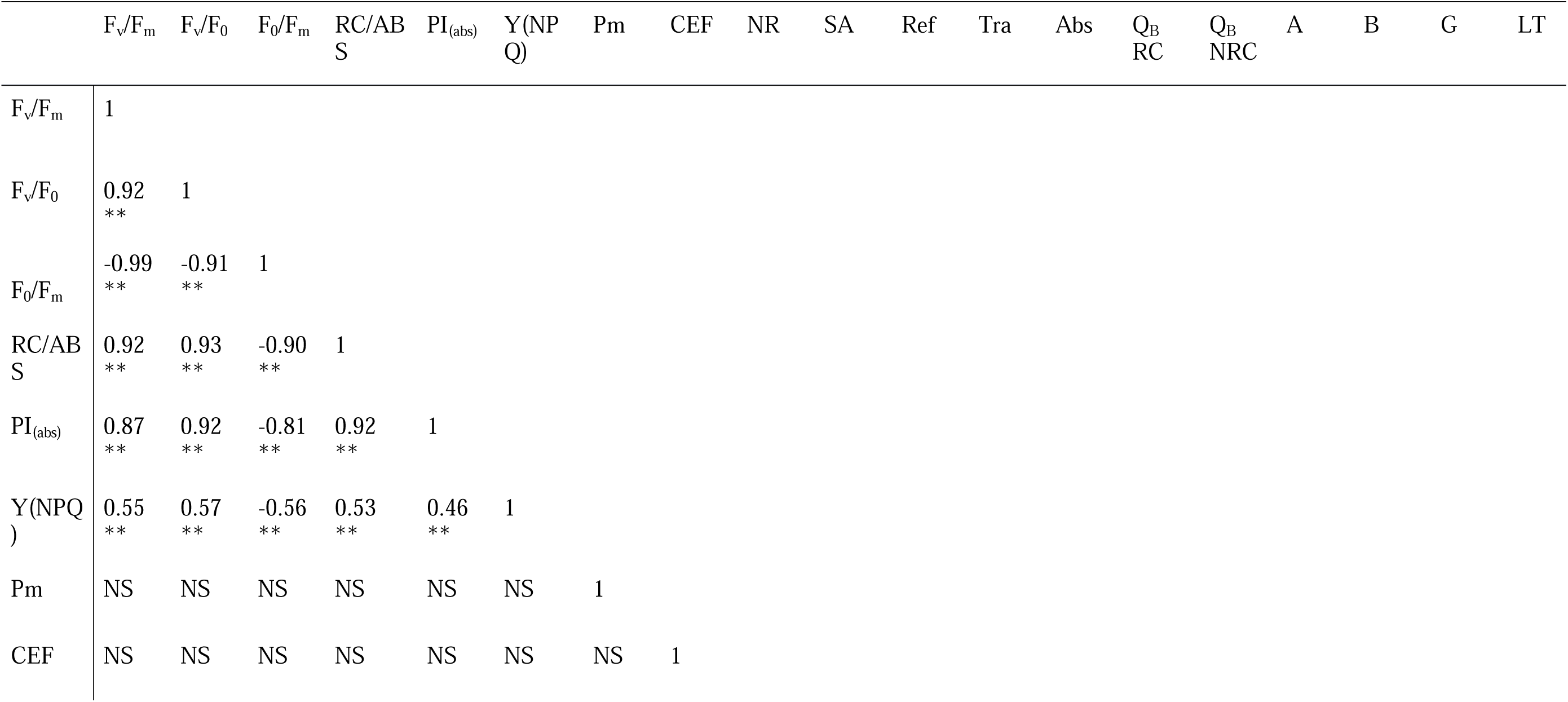
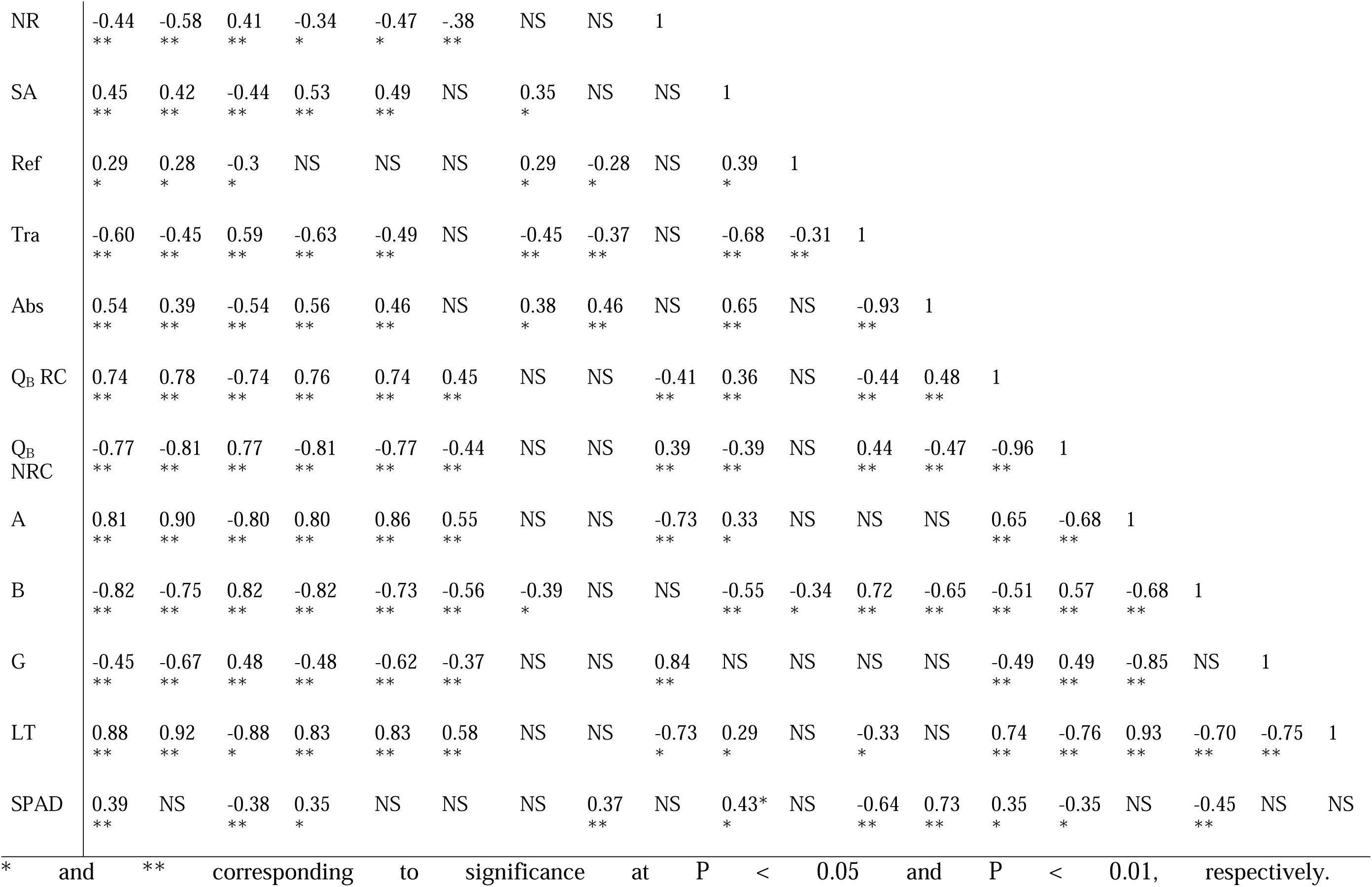
Pearson correlation coefficient between chlorophyll *a* (Chl*a*) fluorescence transients, redox state of P700, cyclic electron flow in PSI, night respiration, stomatal functioning, heterogeneity of PSII, leaf thermal and optical properties and chlorophyll index regardless of mangrove species (*Bruguiera gymnorhiza* [BG] and *Kandelia obovata* [KO]), populations (BGG: BG from Guangxi, China, BGH: BG from Hainan, China, BGI: BG from Iriomote, Japan, BGO: BG from Okinawa, Japan, KOG: KO from Guangxi, China, KOF: KO from Fujian, China, KOI: KO from Iriomote, Japan and KOO: KO from Okinawa, Japan) and temperature conditions (favorable [FDNT] and cold [CDNT; cold wave] day and night temperatures). F_v_/F_m_: maximum quantum yield of PSII in the dark-adjusted state (relative units); F_v_/F_0_: status of water splitting complex (relative units), F_0_/F_m_: thylakoid membrane damage (relative units); RC/ABS: active PSII reaction centers per chlorophyll; PI_(abs)_:performance index; Y(NPQ): effective quantum yield of NPQ; P_m_: maximum photo-oxidizable P700 (ΔI/I); CEF: cyclic electron flow; NR: night respiration; SA: stomatal area; Ref: Reflectance; Tra: transmittance; Abs: absorptance; Q_B_ RC: Q_B_ reducing center; Q_B_ NRC: Q_B_ non-reducing center; A: PSIIα center; B: PSIIβ center; G: PSIIγ center; LT: leaf temperature; SPAD: chlorophyll index.

**Supplementary Table 6.**
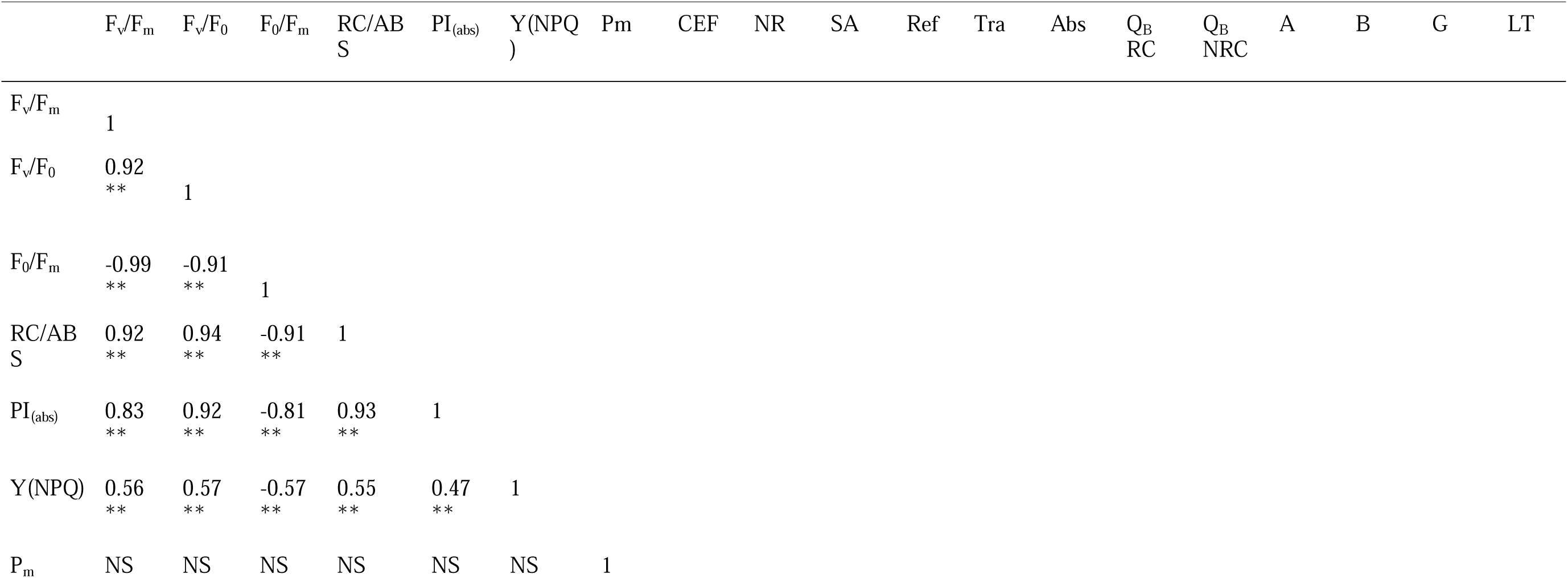
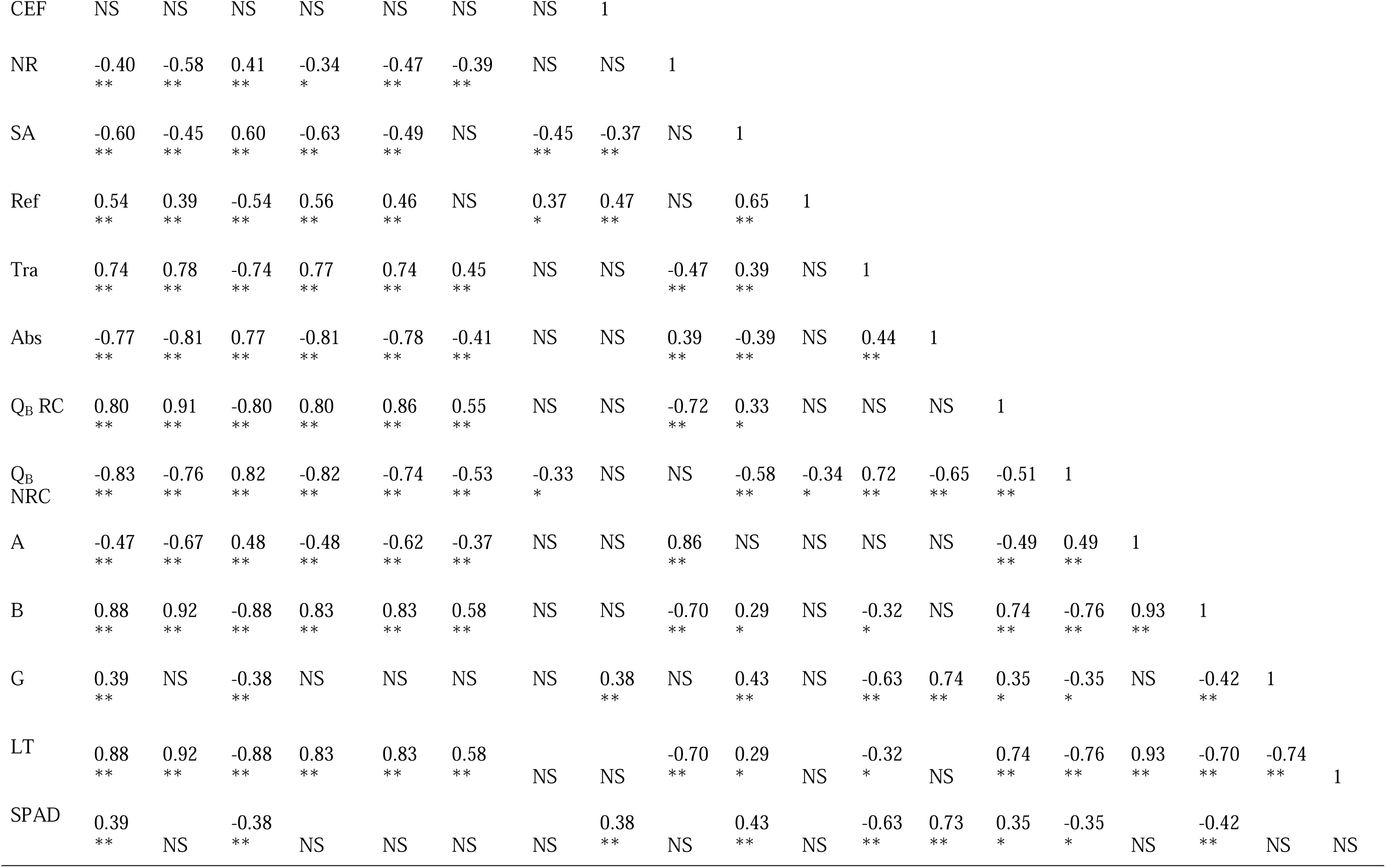
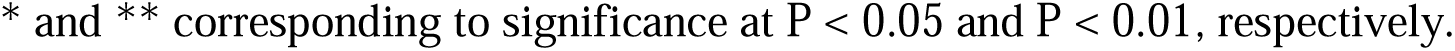
Pearson correlation coefficient between chlorophyll *a* (Chl*a*) fluorescence transients, redox state of P700, cyclic electron flow in PSI, night respiration, stomatal functioning, heterogeneity of PSII, leaf thermal and optical properties and chlorophyll index regardless of mangrove species (*Bruguiera gymnorhiza* [BG] and *Kandelia obovata* [KO]) and its populations (BGG: BG from Guangxi, China, BGH: BG from Hainan, China, BGI: BG from Iriomote, Japan, BGO: BG from Okinawa, Japan, KOG: KO from Guangxi, China, KOF: KO from Fujian, China, KOI: KO from Iriomote, Japan and KOO: KO from Okinawa, Japan) under favorable (FDNT) day and night temperature condition. F_v_/F_m_: maximum quantum yield of PSII in the dark-adjusted state (relative units); F_v_/F_0_: status of water-splitting complex (relative units), F_0_/F_m_: thylakoid membrane damage (relative units); RC/ABS: active PSII reaction centers per chlorophyll; PI_(abs)_:performance index; Y(NPQ): effective quantum yield of NPQ; P_m_: maximum photo-oxidizable P700 (ΔI/I); CEF: cyclic electron flow; NR: night respiration; SA: stomatal area; Ref: Reflectance; Tra: transmittance; Abs: absorptance; Q_B_ RC: Q_B_ reducing center; Q_B_ NRC: Q_B_ non-reducing center; A: PSIIα center; B: PSIIβ center; G: PSIIγ center; LT: leaf temperature; SPAD: chlorophyll index.

**Supplementary Table 7:**
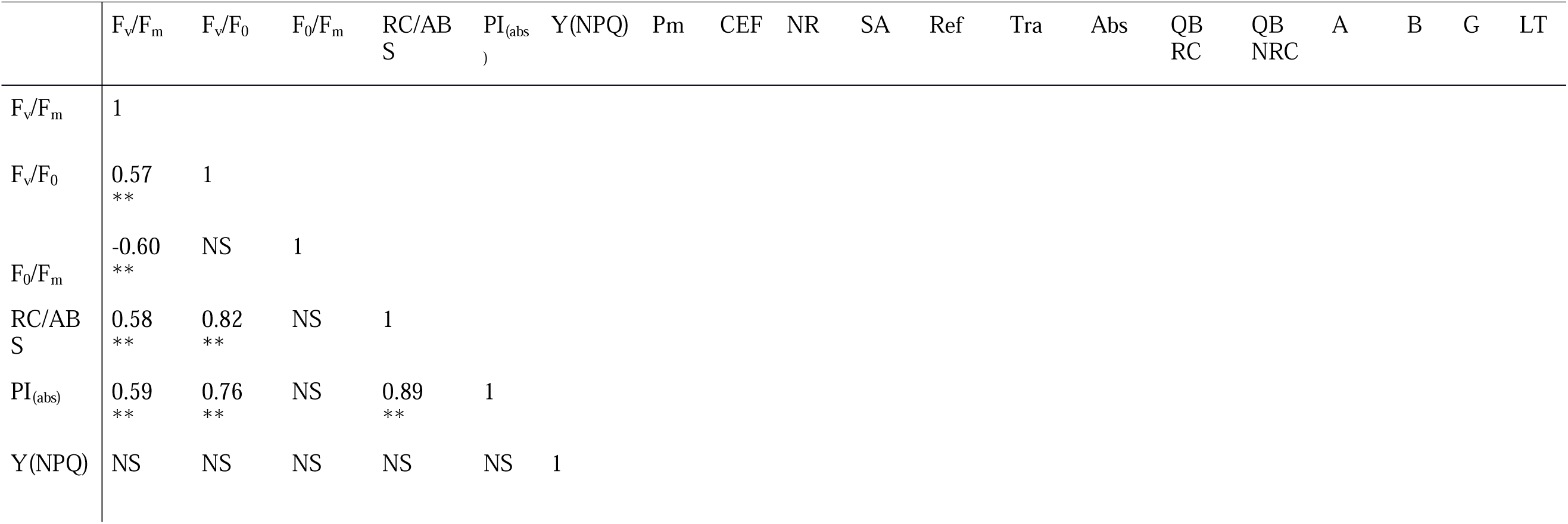
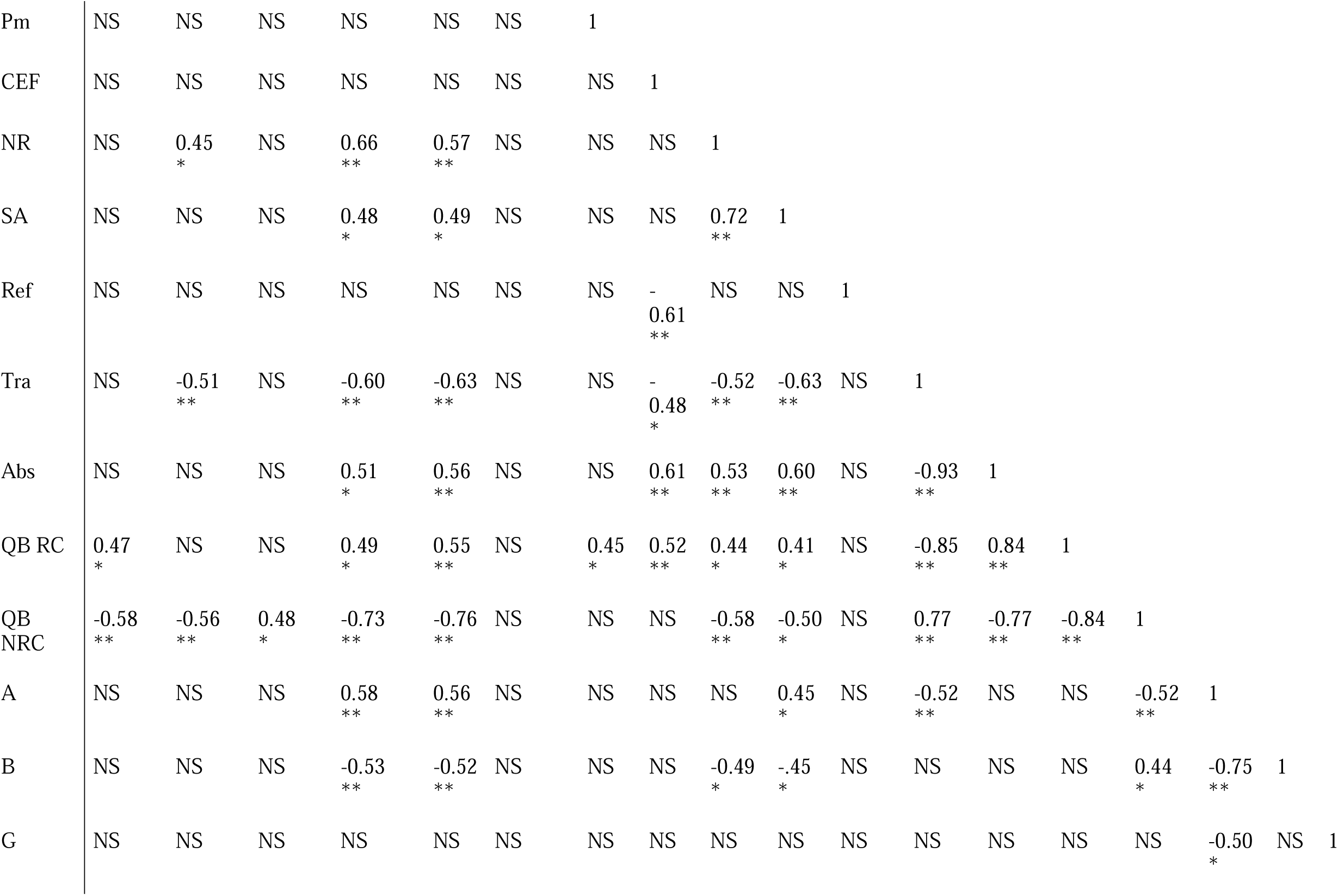
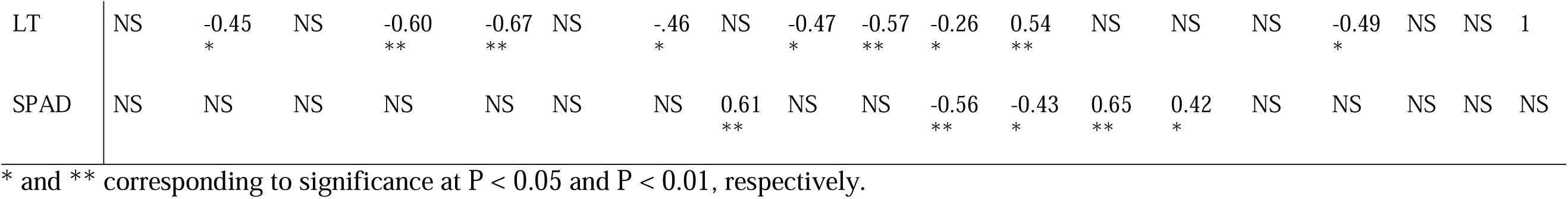
Pearson correlation coefficient between chlorophyll *a* (Chl*a*) fluorescence transients, redox state of P700, cyclic electron flow in PSI, night respiration, stomatal functioning, heterogeneity of PSII, leaf thermal and optical properties and chlorophyll index regardless of mangrove species (*Bruguiera gymnorrhiza* [BG] and *Kandelia obovata* [KO]) and its populations (BGG: BG from Guangxi, China, BGH: BG from Hainan, China, BGI: BG from Iriomote, Japan, BGO: BG from Okinawa, Japan, KOG: KO from Guangxi, China, KOF: KO from Fujian, China, KOI: KO from Iriomote, Japan and KOO: KO from Okinawa, Japan) during cold day and night temperature condition (CDNT; cold wave) day and night temperature condition. F_v_/F_m_: maximum quantum yield of PSII in the dark-adjusted state (relative units); F_v_/F_0_: status of water-splitting complex (relative units), F_0_/F_m_: thylakoid membrane damage (relative units); RC/ABS: active PSII reaction centers per chlorophyll; PI_(abs)_:performance index; Y(NPQ): effective quantum yield of NPQ; P_m_: maximum photo-oxidizable P700 (ΔI/I); CEF: cyclic electron flow; NR: night respiration; SA: stomatal area; Ref: Reflectance; Tra: transmittance; Abs: absorptance; Q_B_ RC: Q_B_ reducing center; Q_B_ NRC: Q_B_ non-reducing center; A: PSIIα center; B: PSIIβ center; G: PSIIγ center; LT: leaf temperature; SPAD: chlorophyll index.

| | $F_v/F_m$ | $F_v/F_0$ | $F_0/F_m$ | RC/ABS | $PI_{(abs)}$ | Y(NPQ) | $P_m$ | CEF | NR | SA | Ref | Tra | Abs | $Q_B$ RC | $Q_B$ NRC | A | B | G | LT |
| --- | --- | --- | --- | --- | --- | --- | --- | --- | --- | --- | --- | --- | --- | --- | --- | --- | --- | --- | --- |
| $F_v/F_m$ | 1 | | | | | | | | | | | | | | | | | | |
| $F_v/F_0$ | 0.57** | 1 | | | | | | | | | | | | | | | | | |
| $F_0/F_m$ | -0.60** | NS | 1 | | | | | | | | | | | | | | | | |
| RC/ABS | 0.58** | 0.82** | NS | 1 |  |  |  |  |  |  |  |  |  |  |  |  |  |  |  |
| $PI_{(abs)}$ | 0.59** | 0.76** | NS | 0.89** | 1 | | | | | | | | | | | | | | |
| Y(NPQ) | NS | NS | NS | NS | NS | 1 |  |  |  |  |  |  |  |  |  |  |  |  |  |

|  |  |  |  |  |  |  |  |  |  |  |  |  |  |  |  |  |  |  |
| --- | --- | --- | --- | --- | --- | --- | --- | --- | --- | --- | --- | --- | --- | --- | --- | --- | --- | --- |
| Pm | NS | NS | NS | NS | NS | NS | 1 |  |  |  |  |  |  |  |  |  |  |  |
| CEF | NS | NS | NS | NS | NS | NS | NS | 1 |  |  |  |  |  |  |  |  |  |  |
| NR | NS | 0.45<br>* | NS | 0.66<br>** | 0.57<br>** | NS | NS | NS | 1 |  |  |  |  |  |  |  |  |  |
| SA | NS | NS | NS | 0.48<br>* | 0.49<br>* | NS | NS | NS | 0.72<br>** | 1 |  |  |  |  |  |  |  |  |
| Ref | NS | NS | NS | NS | NS | NS | NS | -<br>0.61<br>** | NS | NS | 1 |  |  |  |  |  |  |  |
| Tra | NS | -0.51<br>** | NS | -0.60<br>** | -0.63<br>** | NS | NS | -<br>0.48<br>* | -0.52<br>** | -0.63<br>** | NS | 1 |  |  |  |  |  |  |
| Abs | NS | NS | NS | 0.51<br>* | 0.56<br>** | NS | NS | 0.61<br>** | 0.53<br>** | 0.60<br>** | NS | -0.93<br>** | 1 |  |  |  |  |  |
| QB RC | 0.47<br>* | NS | NS | 0.49<br>* | 0.55<br>** | NS | 0.45<br>* | 0.52<br>** | 0.44<br>* | 0.41<br>* | NS | -0.85<br>** | 0.84<br>** | 1 |  |  |  |  |
| QB<br>NRC | -0.58<br>** | -0.56<br>** | 0.48<br>* | -0.73<br>** | -0.76<br>** | NS | NS | NS | -0.58<br>** | -0.50<br>* | NS | 0.77<br>** | -0.77<br>** | -0.84<br>** | 1 |  |  |  |
| A | NS | NS | NS | 0.58<br>** | 0.56<br>** | NS | NS | NS | NS | 0.45<br>* | NS | -0.52<br>** | NS | NS | -0.52<br>** | 1 |  |  |
| B | NS | NS | NS | -0.53<br>** | -0.52<br>** | NS | NS | NS | -0.49<br>* | -0.45<br>* | NS | NS | NS | NS | 0.44<br>* | -0.75<br>** | 1 |  |
| G | NS | NS | NS | NS | NS | NS | NS | NS | NS | NS | NS | NS | NS | NS | NS | -0.50<br>* | NS | 1 |
| LT | NS | -0.45<br>* | NS | -0.60<br>** | -0.67<br>** | NS | -.46<br>* | NS | -0.47<br>* | -0.57<br>** | -0.26<br>* | 0.54<br>** | NS | NS | NS | -0.49<br>* | NS | NS | 1 |
| SPAD | NS | NS | NS | NS | NS | NS | NS | 0.61<br>** | NS | NS | -0.56<br>** | -0.43<br>* | 0.65<br>** | 0.42<br>* | NS | NS | NS | NS | NS |

\* and \*\* corresponding to significance at $P < 0.05$ and $P < 0.01$ , respectively.

**Supplementary Fig. 1:** Representative leaf thermal image of populations of (A and C) *Bruguiera gymnorhiza* (BG) and (B and D) *Kandelia obovata* (KO)] during a cold day and night temperature (CDNT; cold wave). (A) BGI: BG from Iriomote, Japan (B) KOI: KO from Iriomote, Japan, (C) BGH: BG from Hainan, China and (D) KOF: KO from Fujian, China.

**Supplementary Fig. 2:** Microscopic images of real-time status of stomata under favorable day and night temperature (FDNT) [populations of (A) *Bruguiera gymnorhiza* (BG) and (C) *Kandelia obovata* (KO)] and cold day and night temperature (CDNT; cold wave) [populations of (B) *B. gymnorhiza* and (D) *K. obovata*] conditions. BGG: BG from Guangxi, China, BGH: BG from Hainan, China, BGI: BG from Iriomote, Japan, BGO: BG from Okinawa, Japan, KOG: KO from Guangxi, China, KOF: KO from Fujian, China, KOI: KO from Iriomote, Japan and KOO: KO from Okinawa, Japan.

